# Dach1 extends artery networks and protects against cardiac injury

**DOI:** 10.1101/2020.08.07.242164

**Authors:** Brian Raftrey, Ian Williams, Pamela E. Rios Coronado, Andrew H. Chang, Mingming Zhao, Robert Roth, Raquel Racelis, Gaetano D’Amato, Ragini Phansalkar, Karen M. Gonzalez, Yue Zhang, Daniel Bernstein, Kristy Red-Horse

## Abstract

Coronary artery disease (CAD) is the leading cause of death worldwide, but there are currently no available methods to stimulate growth or regeneration of artery networks in diseased hearts. Studying how arteries are built during embryonic development could illuminate strategies for re-building these vessels in the setting of ischemic heart disease. We previously found, using loss-of-function experiments, that the transcription factor *Dach1* is required for coronary artery development in mouse embryos. Here, we report that *Dach1* overexpression in endothelial cells (ECs) extended coronary arteries and improved survival and heart function in adult mice following myocardial infarction (MI). *Dach1* overexpression increased the length and number of arterial end branches, in both heart and retinal vasculature, by causing additional capillary ECs to differentiate into arterial ECs and contribute to growing arteries. Single-cell RNA sequencing (scRNAseq) of ECs undergoing *Dach1*-induced arterial specification indicated that it potentiated normal artery differentiation, rather than functioning as a master regulator of artery cell fate. ScRNAseq also showed that normal arterial differentiation is accompanied by repression of lipid metabolism genes, which were also repressed by Dach1 prior to arterialization. Together, these results demonstrate that increasing the expression level of *Dach1* is a novel pathway for driving specification of artery ECs and extending arterial vessels, which could be explored as a means of increasing artery coverage to mitigate the effects of CAD.

## Introduction

Heart disease is the leading cause of death and is most commonly triggered by atherosclerotic coronary artery disease (CAD)(www.cdc.gov). Although lifesaving interventions do exist, there are significant limitations to current medical and surgical approaches, highlighting the need for novel treatments (Hulot, Ishikawa, & Hajjar, 2016). Given that many forms of heart disease are caused by dysfunctional arteries, one seemingly promising treatment strategy would be to regenerate these diseased coronary arteries.

The coronary blood vascular bed is composed of three vessel subtypes—arterial, venous, and capillary, the latter of which is most numerous and where tissue oxygen exchange occurs. Much research over the last two decades has identified pro-angiogenic proteins, including VEGF and FGF, that primarily stimulate the growth of capillary blood vessels (Giordano et al., 2001). However, clinical trials applying these factors to treat ischemic heart disease have had little success (Cahill, Choudhury, & Riley, 2017; Taimeh, Loughran, Birks, & Bolli, 2013). It has been suggested that a beneficial treatment would also need to grow new arterial vessels in addition to capillary microvasculature (Potente, Gerhardt, & Carmeliet, 2011; Rubanyi, 2013; Schaper, 2009). However, the factors that stimulate the growth of functioning arterial vessels in an established adult vascular bed, such as the heart, are not completely understood (Simons & Eichmann, 2015).

We previously demonstrated that a transcription factor called *Dach1* was important for coronary artery development. EC-specific *Dach1* deletion resulted in a 33% reduction in coronary artery size (Chang et al., 2017). Mechanistically, Dach1 was important for artery development because it potentiated blood flow-guided EC migration into growing arteries. This effect was mediated in part by upregulating expression of the chemokine *Cxcl12* in ECs. These observations are consistent with the known role of Dach1 in other organs and species where it functions to regulate organ size (Davis et al., 2008; Kalousova et al., 2011, Mardon et al., 1994).

While *Dach1* deletion appeared to stunt coronary artery growth by impairing EC migration, it was unclear whether *Dach1* plays additional roles in artery formation. The first step in coronary artery development is the formation of an immature capillary plexus that fills the heart muscle (Kattan, Dettman, & Bristow, 2004; Lavine et al., 2006; Red-Horse, Ueno, Weissman, & Krasnow, 2010). As development proceeds, individual ECs within the capillary plexus differentiate into arterial ECs (Su et al, 2018). These cells, termed pre-artery cells, are intermixed throughout the capillary plexus, but after arterial specification, migrate together to form large arteries. Whether *Dach1* is also involved in these other arteriogenic processes, such as EC specification or mural cell recruitment, is not known.

Here, using gain-of-function experiments, we found that, in addition to its effect on migration, Dach1 also increased artery EC fate specification, in both the developing heart and retinal vasculature. *Dach1* overexpression in plexus and capillary ECs resulted in more pre-artery specification at early stages and additional arterial vessels at later stages. Using scRNAseq, we found that *Dach1* did not widely induce the expression of known artery cell fate determinants, but rather enhanced arterialization in EC subpopulations that are normally receptive to arterial specification. Finally, upregulation of *Dach1* in adult hearts improved survival and cardiac function following experimental MI. Our results indicate that upregulation of *Dach1* could be explored as a target for therapeutically regenerating arterial blood vessels.

## Results

### Dach1 overexpression increases artery specification and arterial branches in the heart

As we found that endothelial *Dach1* was required for coronary arteries to reach their full size during development (Chang et al., 2017), we hypothesized that increasing its expression would promote artery growth. To test this hypothesis, we generated a transgenic mouse that can overexpress *Dach1* in specific cell types at any desired age. A transgene containing the *CAG* promoter upstream of a *Flox-Stop-Flox-Dach1-IRES-EGFP* sequence was inserted into the *ROSA26* locus (Fig. 1A). We next crossed mice carrying the *Dach1^OE^* allele with *ApjCreER*, which is expressed in ECs of the developing coronary capillary plexus and veins but not in differentiated arterial ECs (Chen et al., 2014). Induction of Cre recombinase activity with Tamoxifen in these mice results in excision of the transcriptional stop sequence and permanent co-expression of *Dach1* and EGFP in plexus ECs. We observed high levels of Cre-dependent recombination and *Dach1* expression, which we inferred based on EGFP (Fig. 1B) and anti-Dach1 immunofluorescence (Fig. 1C). In our studies, Tamoxifen was given to experimental (*ApjCreER;Dach1^OE^*) and control (Cre-negative *Dach1^OE^*) mice. However, we also verified that neither transgene affected coronary development in the absence of Tamoxifen (data not shown). This genetic tool was then used to explore the effects of *Dach1* overexpression on artery formation.

**Figure 1.**
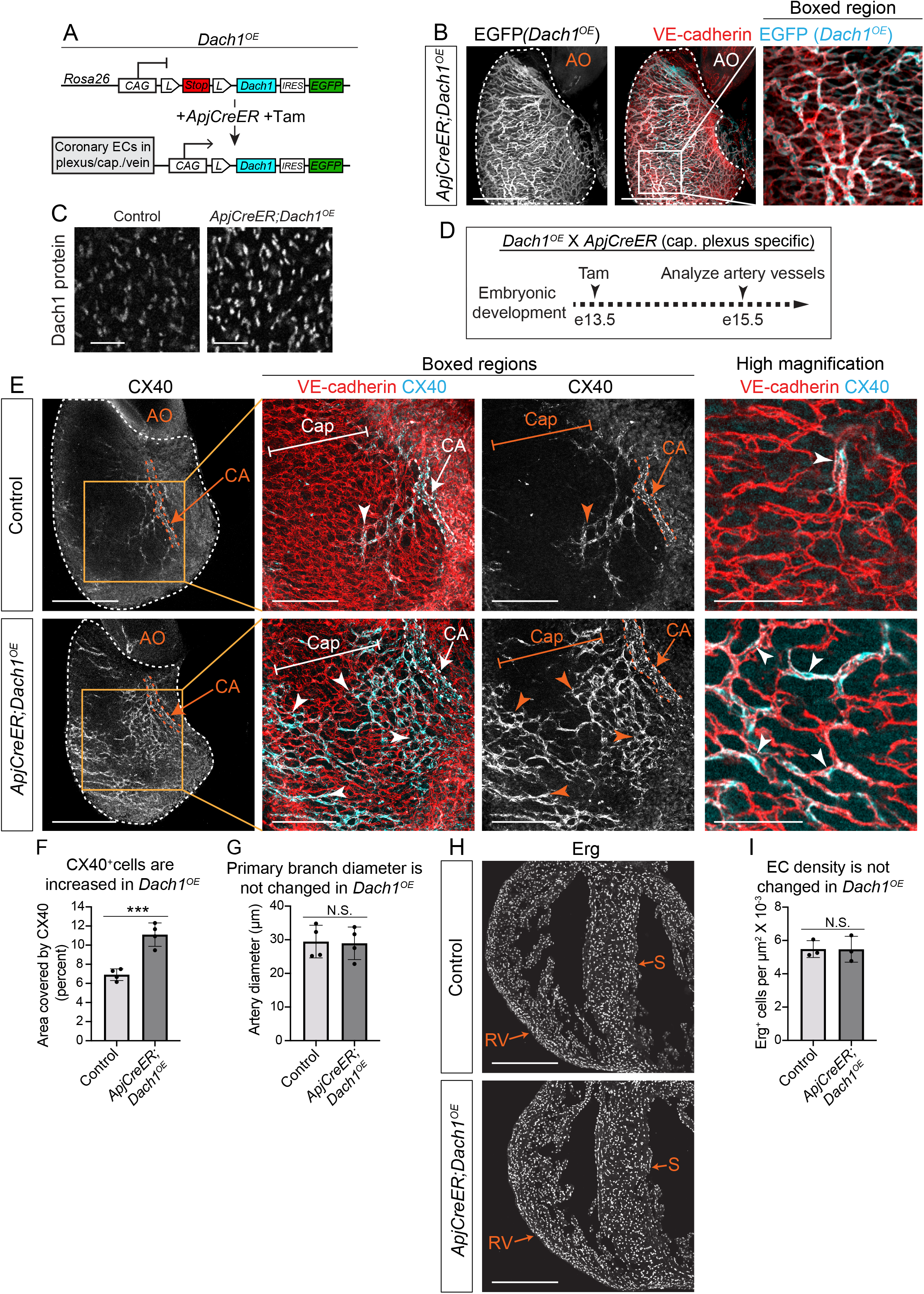
*Dach1^OE^* increases artery specification during coronary artery development. **A**) *Dach1^OE^* transgenic mouse line. **B**) Confocal image of an *ApjCreER;Dach1^OE^* mouse heart (e15.5) immunostained for EGFP to assess recombination rate in the transgenic line. **C**) Dach1 immunostaining in *Dach1^OE^* hearts. **D**) Experimental strategy in **E**-**I**. **E**) E15.5 hearts imaged on the right lateral side show an increase in CX40 staining in *Dach1^OE^* capillaries (arrowheads). **F** and **G**) Quantification of area immunostained by CX40 (**F**) and artery diameters (**G**) in e15.5 hearts (n=4 control, n=4 *Dach1^OE^*). **H**) Erg immunostaining e15.5 heart sections. **I**) Quantification of endothelial density revealed no significant change. (n=3 control, n=3 *Dach1^OE^*) AO= Aorta, CA= Coronary Artery, Cap= Capillary S= Septum, RV= Right Ventricle. Scale bar= 400 μM in B, E (whole heart), and H. Scale bar= 200μM in E (boxed area). Scale bar= 100μM in E highest magnification. Scale bar= 50μM in C. ***=P<.001, N.S.= not significant, all data represent mean+-SD.

We induced *Dach1^OE^* at embryonic day (e) 13.5, when arterial EC differentiation normally begins, and harvested embryos at e15.5 (Fig. 1D). At this time, single, pre-artery ECs that have differentiated within the immature capillary plexus have begun to coalesce into coronary arteries that can be morphologically distinguished (Su et al., 2018). The effect of *Dach1^OE^* on artery development was assessed by immunostaining hearts for ECs (VE-cadherin) and the arterial EC marker Connexin 40 (CX40). CX40-positive artery ECs in control hearts were found lining larger diameter arterial vessels and in the capillary plexus as cells that have been specified to the artery fate but have not yet assembled into the artery, i.e. pre-artery cells (Fig. 1E, upper panels). In *Dach1^OE^* mutant hearts, we observed a dramatic increase in the number of capillary plexus ECs expressing CX40 (Fig. 1E, lower panels). Quantifying the area of each heart covered by CX40 positive ECs revealed a 71% increase in *Dach1^OE^* hearts (Fig. 1F). These results indicate that increasing the levels of *Dach1* stimulates pre-artery specification in ECs within the capillary plexus. The width of the primary coronary artery branch stemming directly from the aorta was not significantly changed (Fig. 1G); heart size (Fig. 1H) and number of ECs within the myocardium (Fig. 1I) were also the same between controls and *Dach1^OE^*. Based on these findings, we concluded that *Dach1* overexpression increased the abundance of arterialized ECs without grossly affecting other aspects of cardiac development.

To investigate whether the increase in pre-artery specification through *Dach1* overexpression affected arterial morphology at later stages, we analyzed CX40 staining at e17.5 when arteries are more mature (Fig. 2A). Immunostaining for CX40 revealed that distal artery branches were more numerous while artery diameters in the main branches were not changed (Fig. 2B). Again, EC coverage did not appear to be grossly affected as assessed by VE-cadherin staining (Fig. 2B). Summing the lengths of all CX40+ vessels revealed a 79% increase in arterial vessel lengths (Fig. 2C) and counting arterial junctions showed a 334% increase in branching (Fig. 2D). The primary artery diameter was not changed (Fig. 2E). This suggests that the increased pre-artery specification observed at e15.5 precedes the development of excessive distal artery branches at e17.5.

**Figure 2.**
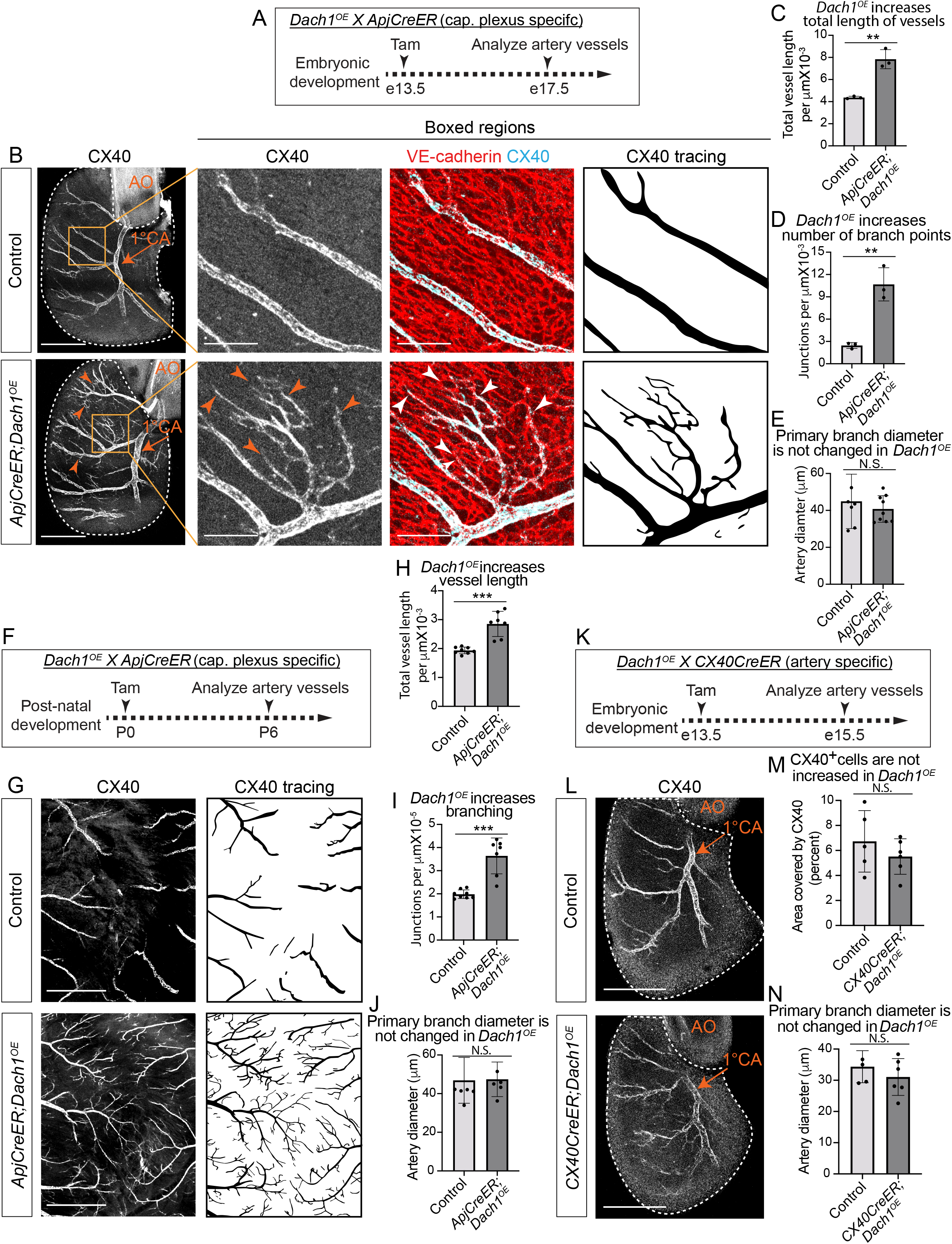
*Dach1^OE^* increases coronary artery branching. **A**) Experimental strategy in **B**-**E**. **B**) Right lateral view of e17.5 hearts. Arrowheads indicate extra artery branches. Boxed regions show the scale of extra CX40^+^ vessels and the normal capillary bed morphology in *Dach1^OE^*. **C**-**E**) Quantification of the total length (**C**) and number of branch points (**D**) in the CX40^+^ vessel network, and primary coronary artery diameters (**E**) in e17.5 hearts (n=7 control, n=9 *Dach1^OE^*). **F**) Experimental strategy in **G**-**J**. **G**) CX40 immunostaining of the ventral surface of post-natal hearts shows increased branching in *Dach1^OE^*. **H** and **I**) Quantification of the total length (**H**) and branch points (**I**) of CX40^+^ vessels, and **J)** measurement of main coronary artery diameters in post-natal hearts (n=7 control, n=6 *Dach1^OE^*). **K**) Experimental strategy to generate artery specific *Dach1^OE^* expression using *Cx40CreER*. **L**) CX40 immunostaining of control and *CX40CreER;Dach1^OE^* hearts. **M** and **N**) Quantification of CX40^+^ area (**M**) and artery diameter (**N**) (n=5 control, n=6 *Dach1^OE^*). AO= Aorta, CA= Coronary Artery. Scale bar= 500μM in G). Scale bar= 400μM in B) (entire heart) and L). Scale bar= 200μM in B) (boxed region).**=p<.01, ***=p<.001, all data represent mean+-SD.

We also induced *Dach1* expression postnatally by dosing nursing mothers with Tamoxifen at postnatal day (P) 0 and analyzing coronary arteries at P6 (Fig. 2F). Similar to embryonic hearts, distal artery branches were more numerous (Fig. 2G, H) with increased branching (Fig. 2I) in the watershed area between the right and left coronary arteries. In contrast, the width of the main coronary artery branch was unchanged (Fig. 2J).

We next sought to determine whether the increase in artery branches resulted from Dach1 activity in capillary plexus ECs or in arterial ECs. In the experiments above, *ApjCreER* will only induce *Dach1^OE^* in the capillary plexus, but its overexpression will be maintained in the arteries that differentiate from these cells. To test whether *Dach1* has the same effects when expressed exclusively in artery ECs, we analyzed e15.5 embryos dosed with Tamoxifen at e13.5 that express *Dach1^OE^* specifically in artery ECs using *CX40CreER* (Fig. 2K). In contrast to *ApjCreER* induction, *CX40CreER;Dach1^OE^* arteries were not significantly different from controls, although there was a trend towards a slight decrease (Fig. 2L-N). Thus, we conclude that *Dach1* increases arterial branching through its activity in capillary plexus ECs.

### Dach1 overexpression increases artery branches in the retina in a cell autonomous manner

We next performed similar experiments in the retina to investigate whether *Dach1*-induced artery specification occurs in other vascular beds. *Dach1* was overexpressed in all ECs using a pan-endothelial Cre, *Cdh5CreER*, and a high dose of Tamoxifen at P0. Then, VE-cadherin and CX40 expression were used to analyze arterial morphology at P7 (Fig. 3A and Supplemental Fig. 1A, B). There was a 2.5 fold increase in the combined length of all CX40-labeled arteries in *Dach1^OE^* retinas (Fig. 3B, C). Interestingly, arterial branches often crossed paths with veins, which does not occur in controls, suggesting a breakdown of arterial-venous repulsion (Fig. 3D, E). These data show that *Dach1* is capable of inducing extension of the arterial network in a vascular bed other than the heart.

**Figure 3.**
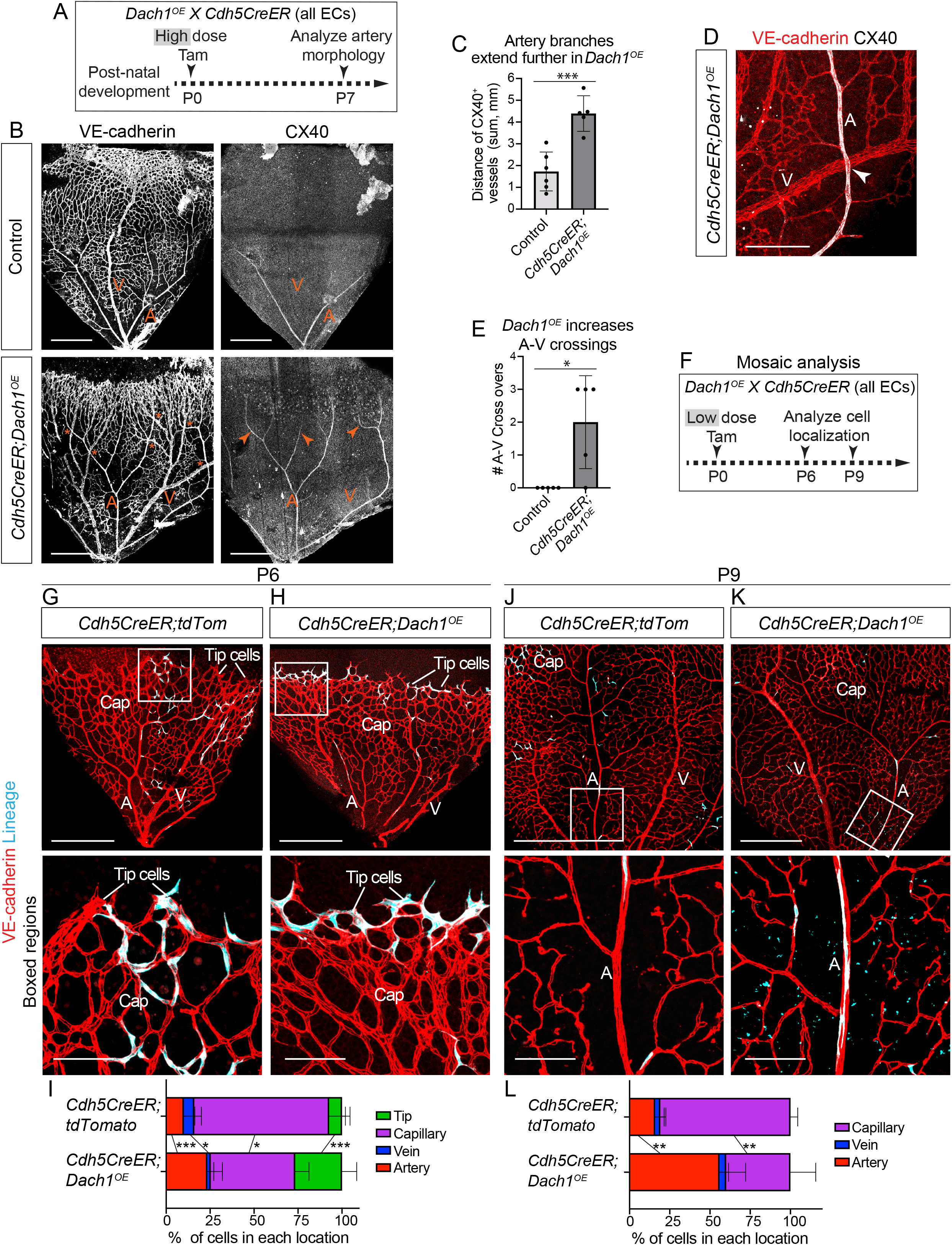
*Dach1^OE^* increases retinal arterialization and promotes endothelial cell migration into arteries. **A**) Dosing strategy for retina vasculature analysis. **B**) Retinas from *Dach1^OE^* pups contained increased number of CX40+ vessel branches (arrowheads) and artery-vein crossing (asterisks). **C**) The total length of all CX40^+^ vessels per retina was greater in *Dach1^OE^* (n=6 control, n=5 *Dach1^OE^*). **D** and **E**) Image (**D**, arrowheads) and quantification (**E**) of artery-vein crossovers in *Dach1^OE^* retinas (n=5 control, n=5 *Dach1^OE^*). **F**) Experimental strategy in **G**-**L**. (**G**-**L**) Images (**G**, **H**, **J**, and **K**) and quantification (**I** and **L**) of control or *Dach1^OE^* cells in retinas from the indicated ages. Boxed regions highlight the tip and capillary cells (**G** and **H**) or artery (**J** and **K**) where there was a differential localization of control and *Dach1^OE^* cells. (**I**: n=5 control, n=7 *Dach1^OE^*; **L**: n=5 control, n=4 *Dach1^OE^*) A= Artery, V= Vein, Cap=Capillary. Scale bar= 400μM in B), G), H), J), and K) (full view). Scale bar= 200μM in D. Scale bar= 100μM in G), H), J), and K) (close up). *=p<.05, **=p<.01 ***=p<.001, all data are mean+-SD.

There were some differences between the heart and retinal vascular beds with respect to *Dach1^OE^*. First, we did not observe any CX40-positive cells outside of arteries within the capillary plexus in either control or *Dach1^OE^* retinas, which is a hallmark of pre-artery cells in the heart. Artery pre-specification has been shown to occur in the retina, but at the tip cell location, i.e. at the migrating front of the growing vasculature (Pitulescu et al., 2017; Xu et al., 2014). However, our data indicate that CX40 does not label pre-specified arterial ECs in the retina as it does in the heart. The second difference was that, in contrast to control hearts, *Dach1^OE^* stunted angiogenesis in the retinas as demonstrated by mildly decreased outward expansion (Fig. 3B). This discordance in phenotypes likely results from the timing of *Dach1^OE^* induction in the two models. In the retina, *Dach1* was overexpressed prior to the initiation of retinal angiogenesis while, in the heart, expression was induced after the coronary plexus was established. Despite these differences, the robust finding is that *Dach1* increased artery branches in both models.

We next investigated whether the arterializing effects of *Dach1* were due to direct cell autonomous activity or were secondary to tissue-level changes which may arise from overexpressing *Dach1* in all retinal ECs. To make this determination, we induced mosaic *Dach1^OE^* at P0 in just 2.3 ± 2.0% of retinal ECs (Fig. 3F) by using a low dose of Tamoxifen. This treatment regimen results in sparse transgene recombination and severely limits the possibility of gross alterations to the retina. *Dach1^OE^* cells are tracked by EGFP expression (Fig. 1A), and *ROSA26;tdTomato* Cre reporter mice were used as a control. Next, the localization of ECs to arteries, capillaries, or veins was determined at P6 and P9. Vessel subtypes were distinguished morphologically using VE-cadherin staining. At P6, the large majority of tdTomato^+^ control cells were within capillary vessels while the remaining were equally distributed among arteries, veins, and tip cells, the latter of which are at the migrating front of the developing vasculature (Fig. 3G). When compared to controls, the number of *Dach1^OE^* cells in arteries and at the tip cell position more than doubled while those in veins decreased (Fig. 3H, I). As pre-specified arterial ECs are known to localize to the tip cell position in the retina (Xu et al., 2014, Pitulescu et al., 2017), the accumulation of *Dach1^OE^* ECs at tips is evidence that it also contributes to arterial pre-specification in the retina, as in the heart. Analyzing cellular distributions three days later at P9 showed that *Dach1^OE^* cells no longer accumulated at tips, but became even more enriched in arteries (Fig. 3J-L). These data demonstrate that *Dach1* overexpression directly causes ECs to follow a path towards arterialization in a cell autonomous fashion.

### Dach1 shifts the endothelial specification trajectory towards arterialization

To understand how *Dach1^OE^* increased EC arterial specification, we performed scRNAseq to identify transcriptomic changes in coronary EC subtypes. Mouse crosses were set up such that in littermates *ApjCreER* induced either *tdTomato* expression (control) or *Dach1^OE^* in plexus ECs at e13.5. Cardiac ECs were then isolated from e15.5 embryos by FACS and processed for scRNAseq using the 10X Genomics platform (Fig. 4A). Analysis of sequencing and alignment parameters confirmed the quality of the data (Supplemental Fig. 2A). Low-quality cells were excluded based on the total number of reads, the number of genes expressed per cell, and the percentage of reads aligned to mitochondrial genes (Supplemental Fig. 2B, C).To focus our analysis of coronary ECs, we also removed the endocardial cell population that lines the lumen of the heart (Supplemental Fig. 2D).

**Figure 4.**
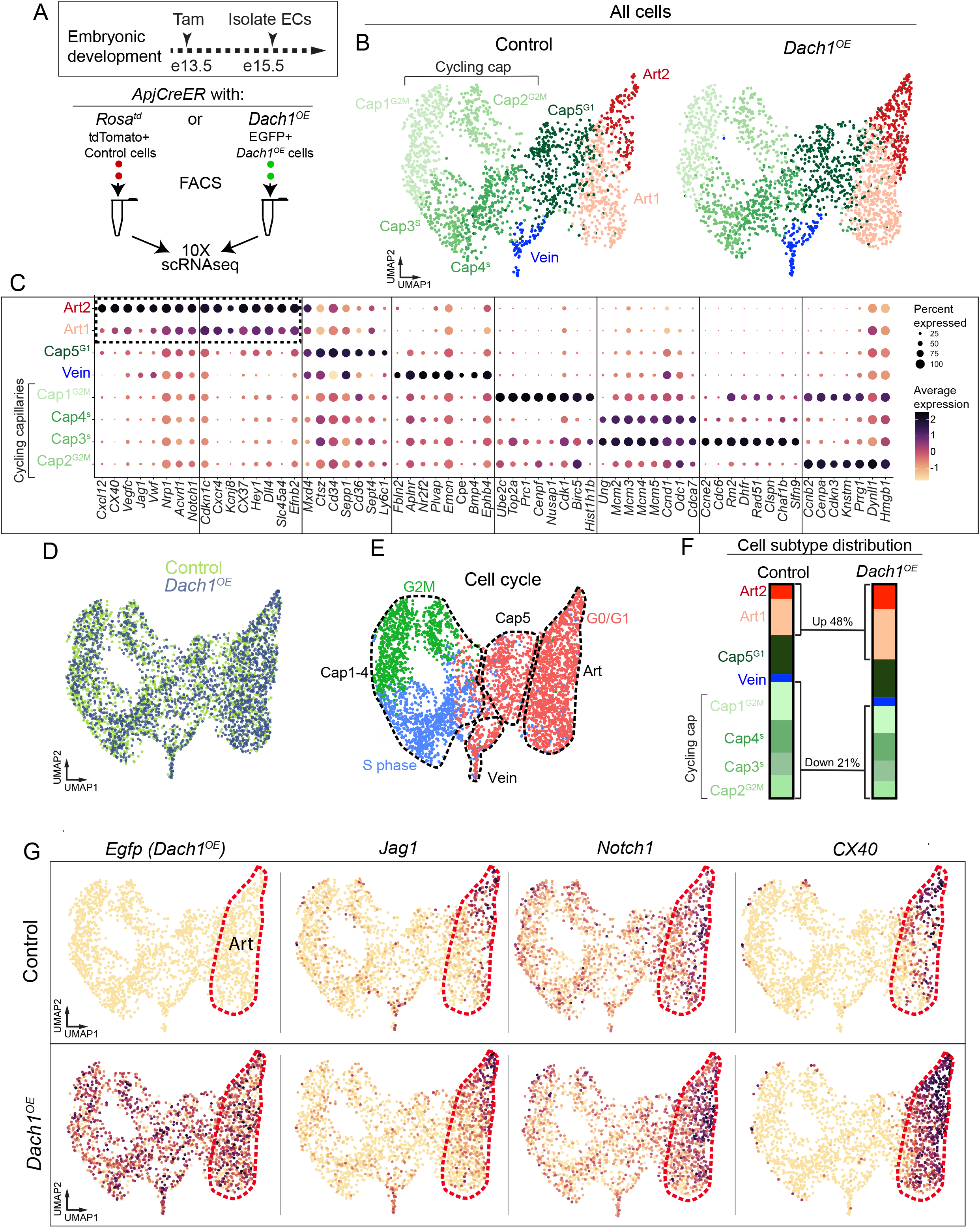
Single cell RNA sequencing of endothelial cells in *Dach1^OE^* hearts. **A**) Littermate e15.5 embryos expressing *ApjCreER* with either *Rosa26tdTomato* or *Rosa26Dach1^OE^* were FACS sorted to isolate coronary endothelial cells for single cell sequencing. **B**) UMAP projections of data showed 8 endothelial cell clusters. **C**) The genes that define each cluster are plotted with their relative expression in each cluster. Boxed region highlights Art1 and Art2 signature genes, which are similar but with increased expression in Art2. **D**) UMAP plot showing that cells from both genotypes overlapped; *Dach1^OE^* did not produce a new subtype. **E**) UMAP plots with both genotypes combined showing cell cycle stage. **F**) Percent of cells in each cluster for both genotype. **G**) Expression of the *Dach1^OE^* transgene and select artery markers.

Control and *Dach1^OE^* datasets were integrated (Stuart et al., 2019) and projected into 2-dimensional space using the uniform manifold approximation and projection (UMAP) algorithm. We then performed unsupervised graph-based clustering to partition ECs into 2 artery (Art1 and Art2), 1 vein, and 5 capillary clusters (Fig. 4B). These EC subpopulations were annotated based on expression of known markers (Fig. 4C). Gene expression profiles suggested that the Art1 cluster was comprised of less mature arterial ECs whereas Art2 arterial ECs were more mature. Specifically, the transcriptomic signature for Art2 consists mostly of increased expression of Art1 markers with the addition of a few mature arterial EC markers such as *Jag1* and *Cxcl12* (Fig. 4C [box] and Supplemental Fig. 3A). All Art1 ECs expressed *Cxcr4* but showed heterogeneous expression of *Cx37* and were negative for *Jag1,* suggesting this cluster is a mix of arterially-skewed capillary ECs and pre-artery ECs (Supplemental Fig. 3A).

Although categorized as distinct clusters, the topography of the UMAP projection and gene expression patterns indicated that EC subtypes exist along a continuum, rather than as completely distinct states (Supplemental Fig. 3B). This phenomenon was particularly evident when projecting cells on an axis of arterial-venous identity (Supplemental Fig. 3C). This is consistent with previous analyses showing that brain and heart capillary ECs exist along a continuum of arterial-venous identity, even in adults (Chen et al., 2020; Su et al., 2018; Vanlandewijck et al., 2018). Nonetheless, cells from all clusters were found in both genotypes (Fig. 4B, D), suggesting that *Dach1^OE^* does not create a new cell identity or transcriptional state not normally present.

We also observed that cell cycle phase, inferred from the enrichment of phase-specific genes, was a major source of transcriptomic variability in ECs. Most capillary ECs (clusters Cap1-4) were either in S or G2/M phase (i.e., cycling) while one capillary cluster (Cap5) and the artery and vein clusters were in G0/G1 (i.e. non-cycling; Fig. 4E). In summary, scRNAseq allowed us to isolate artery, vein, and capillary (cycling and non-cycling) ECs from control and *Dach1^OE^* hearts for subsequent transcriptomic analysis.

Although all EC subtypes were found in both genotypes, *Dach1^OE^* altered the relative number of ECs within specific clusters. *Dach1^OE^* induced a 48% increase in the fraction of arterial ECs while decreasing the fraction of cycling capillary ECs by 21% (Fig. 4F). These findings corroborate our observation of increased CX40^+^ arterial ECs in developing hearts. Importantly, they further demonstrate that these ECs are not only CX40^+^, but also express a full arterial transcriptomic program.

We next sought to uncover the cellular mechanism by which *Dach1^OE^* increased arterial EC specification of capillary ECs. One hypothesis is that Dach1 induces the expression of arterial EC fate determinants in all capillary ECs. Notch transcription factors are the most well recognized artery cell fate determinants described to date (Fang & Hirschi, 2019). In examining differentially expressed genes (DEGs; LogFC > 0.25) between *Dach1^OE^* and control in each cluster, we found that Dach1 does not broadly change the level of Notch pathway genes, or any other validated artery marker, in capillary subpopulations even though it was overexpressed in virtually all ECs (Fig. 4G and Supplemental Fig. 3B). Instead, Dach1 only increased some of these genes in select clusters within our data, including arterial ECs (Fig. 4G and Supplemental Fig. 3B). This observation suggests that Dach1’s arterializing capability is restricted to certain EC subtypes and/or involves upregulating previously uncharacterized arterial fate determinants.

To determine which EC subtypes are sensitive to *Dach1^OE^*-induced arterialization, we calculated artery scores, which were determined by measuring each cell’s enrichment of the genes that defined Art2 in controls (Supplemental Table 1). Consistent with ECs existing along an arterial-venous continuum, control cells were distributed along the entire artery score spectrum (Fig. 5A). The distribution of artery scores in *Dach1^OE^* ECs, on the other hand, was shifted towards higher artery scores (Fig. 5A). This is most likely attributable to elevated artery scores in the artery clusters, as artery scores did not significantly change in other clusters (Fig. 5B and Supplemental Fig. 4A). This is consistent with the specific expansion of *Cx40* in Art1 that is shown in Fig. 4G. These data indicate that Dach1 enhances the arterial transcriptional profile specifically in arterial skewed capillary, pre-artery, and arterial ECs.

**Figure 5.**
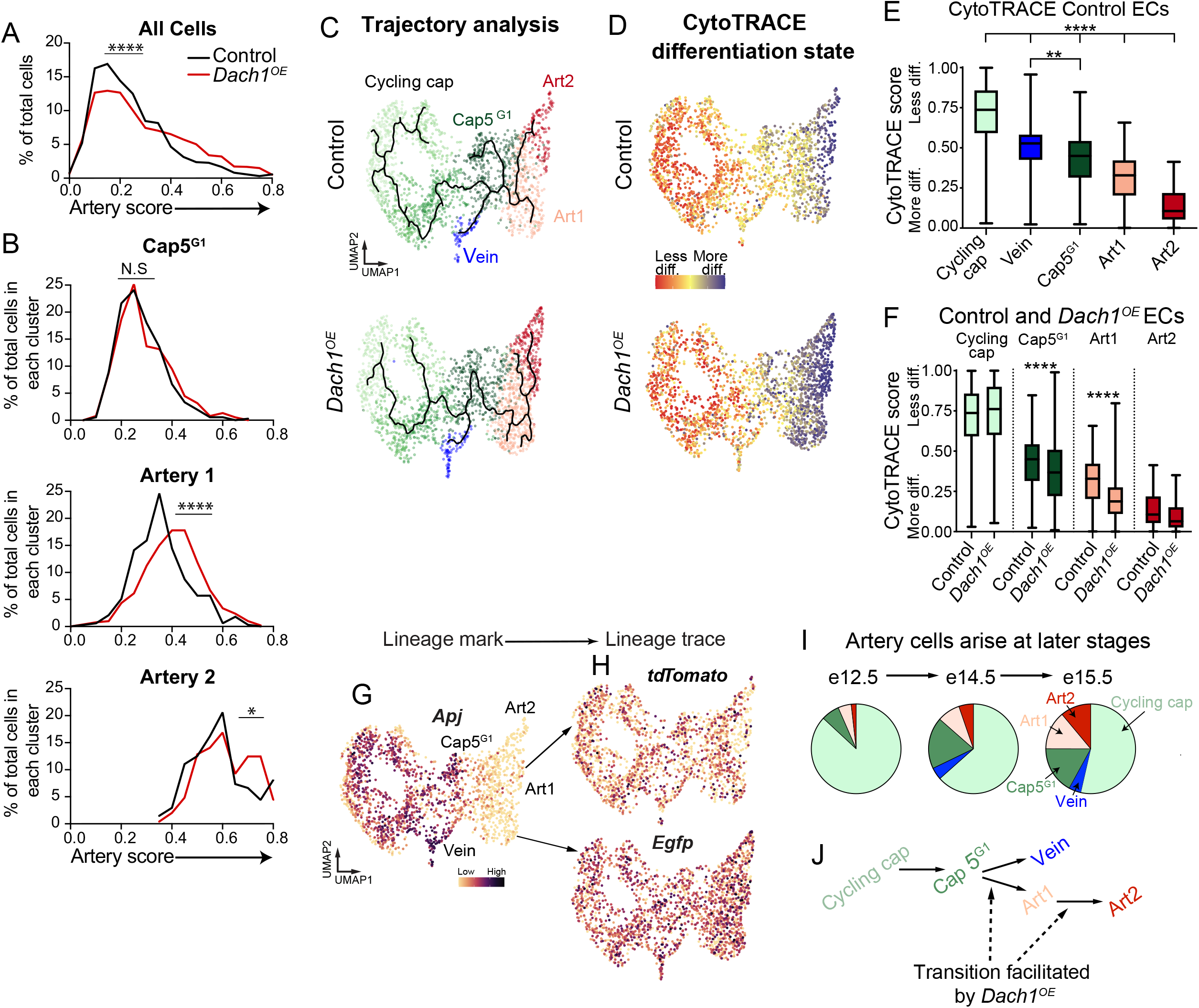
Dach1 enhances artery specification in endothelial cells. **A**) Calculating an artery score for each cell revealed that *Dach1^OE^* shifted the total distribution of cells to a higher artery score. **B**) Within each individual cluster, only the Art1 and Art2 clusters had significant shifts in artery score. **C)** Cell trajectories (solid line) from either genotype inferred using Monocle3. **D**) CytoTRACE differentiation scores for each cell displayed on the UMAP plot. **E**) CytoTRACE scored calculated for controls cells. **F**) CytoTRACE scores were lower (more differentiated) for Cap5^G1^ and Art1in *Dach1^OE^*. **G**) *Apj*, the enhancer/promoter used to drive Cre, is expressed in all endothelial cells except arteries. **H**) The lineage traces from *ApjCreER* (*tdTomato* or *Dach1^OE^-EGFP*) are later expressed in the arterial endothelial cells. **I**) Graphs showing the percent of cells in each indicated cluster in analogous coronary endothelial cell scRNAseq datasets from indicated embryonic days (e). **J**) Model for cell differentiation trajectory during coronary artery development and the proposed influence of Dach1. *=p<.05, **=p<.01 ***=p<.001, ****=p<.0001, box plots are mean+-SD.

In order to determine the effects of *Dach1^OE^* on the trajectory of arterial specification, we first needed to delineate this trajectory in control ECs. Trajectory analysis (Monocle 3) showed that Cycling cap, Cap5^G1^, Art1, and Art2 clusters were transcriptomically connected in series while veins branched off of Cap5^G1^ (Fig. 5C, top panel). We then ordered this lineage connection by developmental stage using CytoTRACE (Cellular Trajectory Reconstruction Analysis using gene Counts and Expression). This method predicts cellular differentiation potential by calculating the expression of genes that correlate with the number of genes expressed per cell (Gulati et al., 2020). CytoTRACE scored the Cycling cap cluster as the least differentiated followed by Vein, Cap5^G1^, Art1, and then finally Art2 as the most differentiated (Fig. 5D, top panel, and E).

Because the above data were collected from a single time point, we sought additional evidence for this developmental trajectory. First, we determined the pattern of lineage tracing labels among different EC subtypes. These data showed that *Apj*, the promoter that drives Cre-mediated *tdTomato* (control) and *EGFP* (*Dach1^OE^*) expression, is only expressed in Cycling cap, Cap5^G1^, and Vein ECs (Fig. 5G and Supplemental Fig. 4B). Thus, these are the only EC subtypes that would initially express *tdTomato* or *EGFP* following Cre induction at e13.5. By e15.5, *tdTomato* (control) and *EGFP* (*Dach1^OE^*) lineage expression was seen in arterial ECs, indicating that the artery clusters developed from *Apj*-positive capillary and/or vein ECs (Fig. 5H). We also integrated this e15.5 *Apj*-traced EC dataset with e12.5 and e14.5 *Apj-*traced EC datasets that we had described previously (Su et al., 2018). Quantifying the ratio of cells at each time point showed that higher proportions of Art1 (less mature) precede higher Art2 (more mature) proportions (Fig. 5I). Considering all of these findings, we propose that the differentiation trajectory of arterializing ECs involves the following steps: 1) exit of proliferating capillary ECs from the cell cycle and differentiation to a non-cycling capillary subtype (Cap5^G1^), 2) initial pre-artery/artery specification (Art1), and 3) full differentiation into mature arterial ECs (Art2)(Fig. 5J).

We next determined how *Dach1^OE^* influences this developmental progression. Trajectory analyses revealed that Cap5^G1^ ECs in *Dach1^OE^* are more linearly connected with arterial ECs (Fig. 5C, lower panel). *Dach1^OE^* Cap5^G1^ and Art1 cells were significantly more differentiated as scored by CytoTRACE (Fig. 5D, lower panel, and F) while the differentiation state of Cycling cap, Vein, and Art2 ECs were not changed (Fig. 5D and F). These data support a model where Dach1 is not an arterial cell fate determinant per se, but rather potentiates differentiation and arterialization in receptive cells such as Cap5^G1^ and the pre-artery cells in Art1 (Fig. 5J).

### Gene expression changes in Dach1^OE^

To investigate whether *Dach1^OE^* might regulate arterial genes not previously known to play a role in arterialization, we compared DEGs between *Dach1^OE^* and controls with genes that were positively or negatively enriched in control artery ECs in our dataset. Each cluster had between 50-160 DEGs, including both upregulated and downregulated genes (Fig. 6A). 38% of the Vein and 34% of the Cycling cap DEGs that were upregulated in *Dach1^OE^* were on the list of 498 genes whose increase defined the Artery 2 cluster in control hearts, i.e. “arterial EC genes” (Fig. 6B, Supplementary table 1). More than 50% of the upregulated DEGs in *Dach1^OE^* Cap5^G1^, Art 1, and Art 2 were also arterial EC genes (Fig. 6B). In contrast, a much smaller percentage of the DEGs upregulated in *Dach1^OE^* were on the list of 436 that were downregulated in control arterial ECs, i.e. “non-arterial EC genes” (Fig. 6B, and Supplementary table 1). When considering genes downregulated by *Dach1^OE^*, there were much lower percentages of overlap with arterial EC genes and non-arterial EC genes, and there was no differential pattern between the two types (Fig. 6C). These patterns suggest that all *Dach1^OE^* ECs may contain some level of priming towards an arterial fate through the induction of arterial genes not previously correlated with artery cell fate specification, and that there is a stronger effect on induction of artery genes rather than repression of non-artery genes.

**Figure 6.**
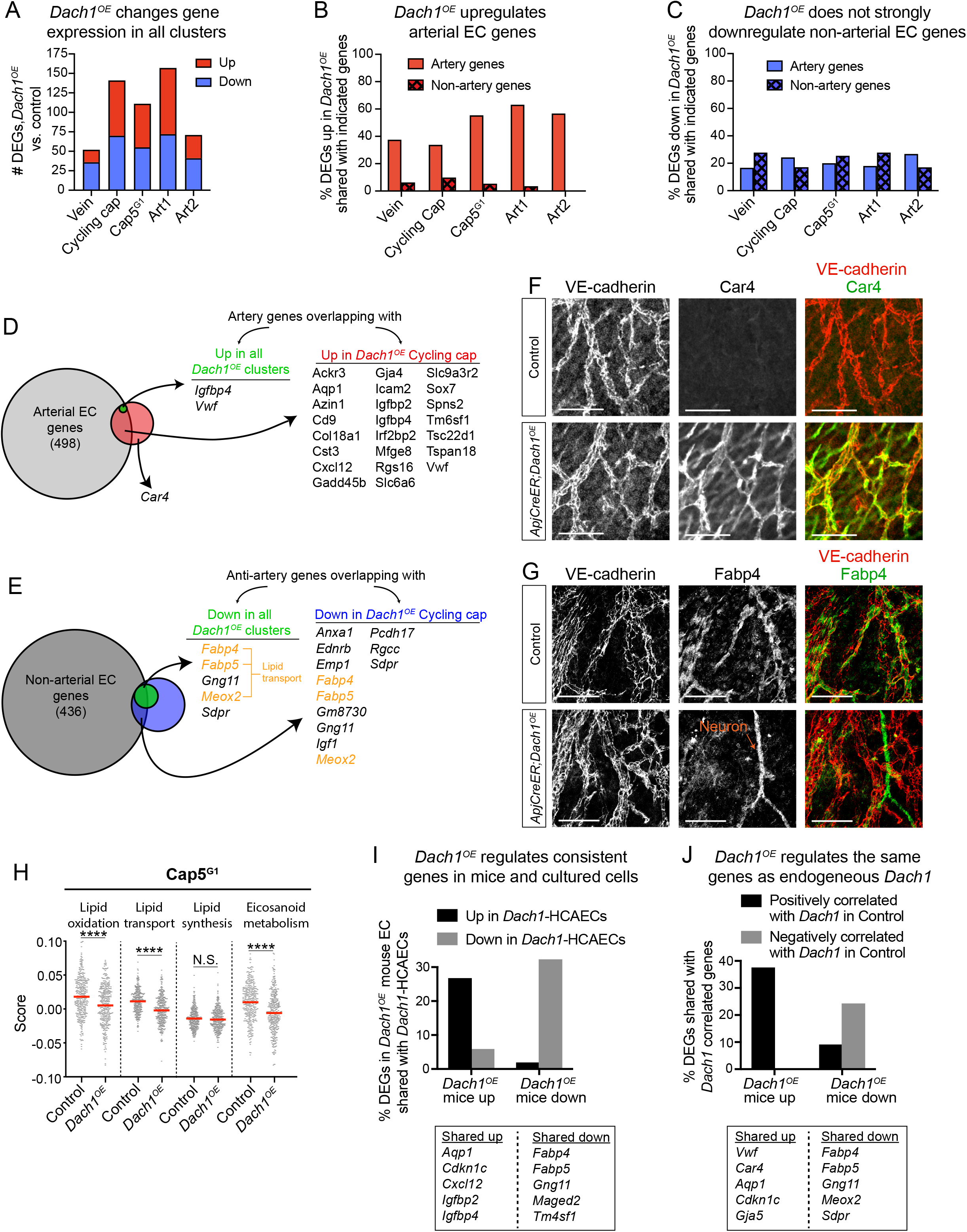
Specific gene expression changes in *Dach1^OE^*. **A**) The number of differentially expressed genes (DEGs) when separately comparing control and *Dach1^OE^* cells from each cluster. **B** and **C**) Genes that are either positively or negatively enriched in control artery clusters we identified and termed “artery genes” and “non-artery genes”, respectively. DEGs that were either up- or down-regulated by *Dach1^OE^* in each cluster were then compared to these lists. Upregulated DEGs in *Dach1^OE^* have strong overlap with artery genes (**B**) while downregulated DEGs in *Dach1^OE^* have less overlap with non-artery genes (**C**). **D** and **E**) Venn diagrams showing overlap of artery (**D**) and non-artery (**E**) genes with DEGs either up- or down-regulated by *Dach1^OE^* in all clusters or cycling capillaries, respectively. **F** and **G**) Validation of scRNAseq genes using immunofluorescence on e15.5 hearts. *Dach1^OE^* increased *Car4* (**F**) and decreased *Fabp4* (**G**) in coronary endothelial cells. **H**) Scores for lipid pathways show a reduction in score for Cap5^G1^ in *Dach1^OE^*. **I**) DEGs shared between endothelial cells experiencing *Dach1* overexpression in either developing mouse hearts (*Dach1^OE^*) or primary cell culture (*Dach1*-HCAECs). **J**) Overlap between mouse *Dach1^OE^* DEGs and genes that are positively or negatively correlated with endogenous *Dach1* in control hearts. Scale bars= 100μM, ****=p<.0001, N.S = not significant.

We next sought to find genes that might be directly regulated by Dach1. We reasoned that such genes would be independent of EC subtype and, therefore, differentially expressed in all clusters. A universal DEG list contained 2 upregulated and 13 downregulated genes that were changed in all clusters. Both upregulated genes (*Igfbp4* and *Vwf*) were arterial EC genes (Fig. 6D). Of the universally downregulated genes, 7 were non-arterial EC genes, and 2 of those— *Fabp5* and *Sdpr—*were Vein genes (Fig. 6E). Interestingly, two of the downregulated genes were involved in lipid transport, and a third, Meox2, is a transcription factor that regulates the expression of those two lipid transporters (Fig. 6E, yellow)(Coppiello et al., 2015). A subset of these DEGs were validated using immunofluorescence (Fig. 6F and G). We next investigated additional lipid metabolism and signaling pathway genes and found that genes associated with lipid transport, lipid oxidation and eicosanoid metabolism were significantly downregulated by *Dach1^OE^* (Fig. 6H and Supplemental Fig. 5A-D). These pathways are not typically linked to artery development, but suggest that suppression of lipid metabolism might be involved. In summary, universal DEGs overlapped with arterial EC and non-arterial EC genes in our dataset, but did not include known mediators of arterial EC fate determination, suggesting that if *Dach1^OE^* primes all ECs, it is through hitherto unrecognized arterial pathways (such as suppression of lipid metabolism).

To more robustly identify genes regulated directly by *Dach1*, *Dach1^OE^* scRNAseq DEG lists were cross-referenced with a previously generated bulk RNAseq dataset from cultured human coronary artery endothelial cells that overexpressed *Dach1* (*Dach1*-HCAECs) (Chang et al., 2017). 26.4% of the genes upregulated in *Dach1^OE^* mouse coronary ECs were also upregulated in *Dach1*-HCAECs, and 30.7% of the downregulated genes overlapped with those downregulated in cultured cells (Fig. 6I). Notable genes regulated in both experimental systems were *Cxcl12, Aqp1, Cdkn1c,* and some of the aforementioned lipid transport genes (Fig. 6I). Therefore, overexpression of *Dach1* activates a consistent set of downstream genes in ECs.

Next, we investigated whether overexpression of Dach1 through the *Dach1^OE^* transgene reflects the set of genes that are endogenously regulated by *Dach1*. We analyzed cells from control hearts only, and correlated the expression of endogenous *Dach1* to the expression of all other genes. We selected the top 500 genes that were positively and negatively correlated with endogenous *Dach1* and compared those to the DEGs from *Dach1^OE^*. This revealed a strong overlap of the DEGs either up or down with *Dach1^OE^* in the list of genes positively or negatively correlated with endogenous *Dach1* expression, respectively (Fig. 6J). Prominent on these lists were arterial EC and non-arterial EC genes, such as *Aqp1* and lipid transporters, respectively. This supports the idea that overexpressing *Dach1* affects similar transcriptional programs as the endogenous *Dach1*, and that this transcription factor regulates artery and lipid transport genes in multiple settings.

Since arterial specification has been linked to cell cycle exit (Fang et al., 2017; Su et al., 2018), we noted that *Cdkn1c* expression, a cell cycle inhibitor, was associated with Dach1 in multiple scenarios (Fig. 6I and J). We performed EdU incorporation experiments aimed at assessing whether Dach1 decreased cell cycling. These were focused on non-arterial regions of the heart to avoid confounding results due to cell cycle exit being linked to arterial differentiation. EdU incorporation was not different with *Dach1^OE^* in vessels on the surface of the heart where arterial differentiation does not occur (Supplemental Fig. 5E). It was decreased within intramyocardial regions, but this could be due to the increased arterial differentiation at that location (Supplemental Fig. 5E, F). EdU incorporation was also not different in the retina (Supplemental Fig. 5G). Thus, although not affecting cell cycling in all cells, upregulation of *Cdkn1c*, could potentiate cell cycle arrest in the presence of other artery differentiation signals.

### Dach1 overexpression supports recovery from MI

Following our observations that *Dach1* overexpression extended the arterial vasculature, we investigated whether it could also improve outcomes following experimental myocardial infarction (MI). To test this, we permanently ligated the left anterior descending coronary artery of 12-week-old adult mice with *ApjCreER*-induced *Dach1* overexpression in capillary and venous ECs for 6 weeks prior to injury (Fig. 7A). Quantification showed that this strategy resulted in recombination, i.e. *Dach1* overexpression, in 64.5±14% of coronary ECs (Fig. 7B).

**Figure 7.**
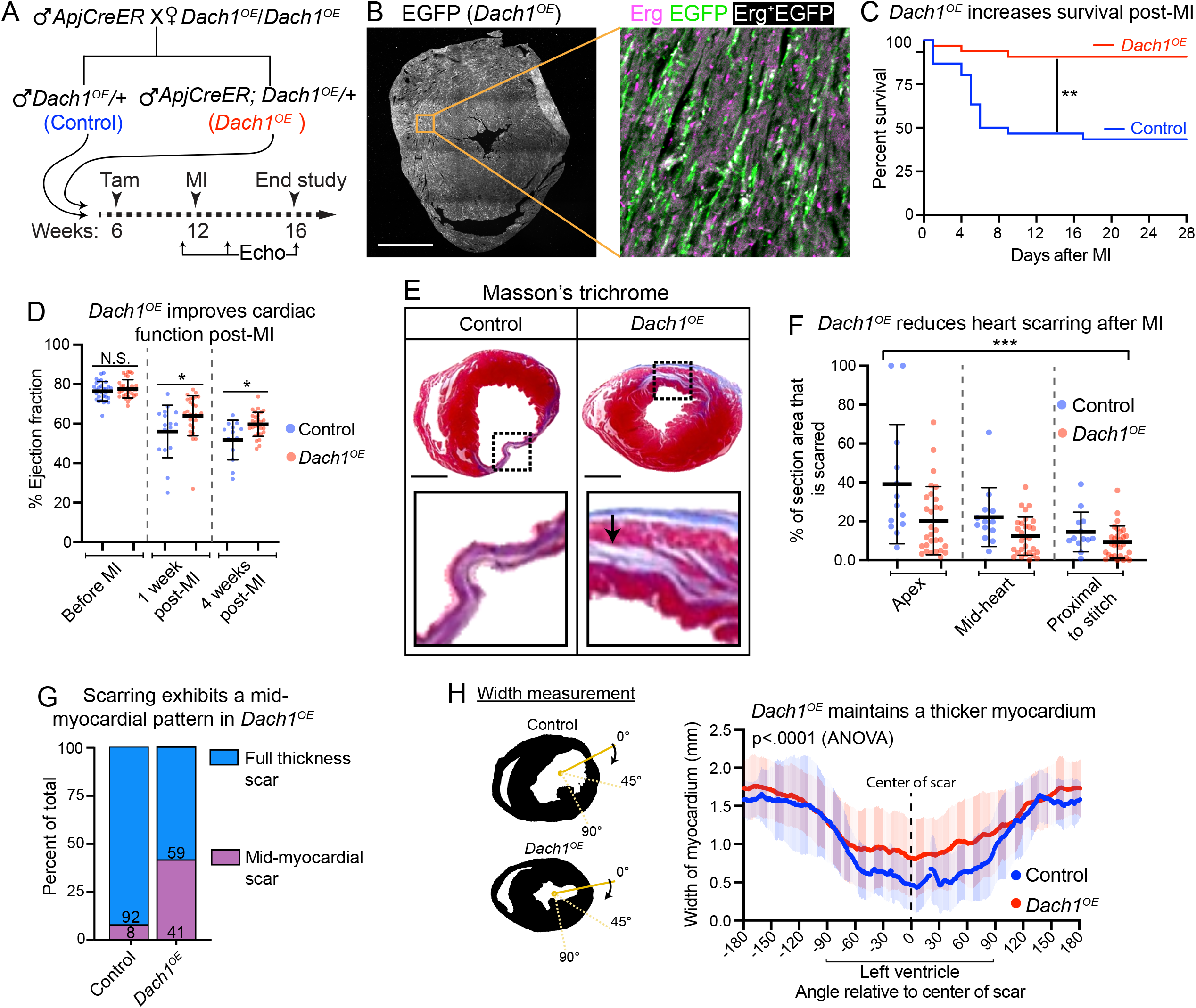
*Dach1* overexpression promotes survival after myocardial infarction. **A**) Experimental strategy for myocardial infarction (MI) study. **B**) Recombination of the *Dach1^OE^* allele in adult hearts following the Tamoxifen dosing strategy in (**A)**. **C)** Survival curve during 4-weeks post-MI. (n=30 control, n=32 *Dach1^OE^*) **D**) Percent ejection fraction at the indicated time points (n=30 control, n=32 *Dach1^OE^*). **E**) Hematoxylin & Eosin staining on representative hearts. **F)** Percent of total myocardium stained with Masson’s Trichrome in sections from three levels posterior to the ligation (n=13 control, n=29 *Dach1^OE^*). **G)** Quantification of scarring pattern. Arrows in **(E)** highlight an example of a mid-myocardial scar. **H)** The width of the myocardium at 360 angles around the heart. In the left ventricle where the infarct was induced, *Dach1^OE^* better preserved myocardial thickness when compared to controls. Lines are average of n=13 control and n=29 *Dach1^OE^* while shading indicates S.D. *=p<.05, **=p<.01, ***=p<.001, N.S = not significant. Error bars show mean+-SD. Log-rank test was used in (**A)**, t-test in (**B)**, Two way ANOVA in (**F)**, (**H)**. Scale bar=1mm in (**B)**, (**E)**.

Experimental MI resulted in 43.3% survival for control animals (Fig. 7C). Most fatalities occurred within the first 10 days post-MI. In contrast, *Dach1^OE^* mice exhibited an 90.6% survival rate (Fig. 7C). Echocardiography post-MI showed that *Dach1^OE^* mice had a significantly higher ejection fraction than controls at both weeks 1 and 4 (Fig. 7D). A control experiment using the same genotypes without Tamoxifen injection showed that this rescue effect was not due to presence of the *ApjCreER* transgene (data not shown).

Histological analysis at 4 weeks revealed a reduction in fibrosis in *Dach1^OE^* compared with controls (Fig. 7E and F). In 92% of WT mice, fibrotic scars extended the full thickness of the myocardium, whereas this pattern was seen in only 59% of *Dach1^OE^* hearts (Fig. 7G). In the hearts that did not have a full thickness scar, fibrosis was limited to the mid-myocardial region (Fig. 7E and G). Measuring myocardial thickness at sequential points around the entire heart revealed an increase in thickness within the left ventricle in *Dach1^OE^* hearts (Fig. 7H).

These results demonstrate that overexpression of *Dach1* in coronary ECs improves overall survival, cardiac function, and myocardial scarring post-MI. These observations and the site of overexpression are consistent with *Dach1* inducing vascular changes that provide protective blood flow in the face of coronary occlusion. This could be due to the arterialization effects of Dach1 (Fig. 1–5) or other noted changes in gene expression (Fig. 6). A detailed analysis of the underlying cause of cardioprotection will be the subject of future studies.

## Discussion

Coronary artery ECs differentiate from capillary precursor cells; however, the transcriptional regulators of this capillary-to-artery transition remain incompletely discovered. Here, we found that overexpression of the transcription factor *Dach1* drove ectopic arterial EC specification in capillary ECs of the coronary vasculature. These extra arterial ECs contributed to artery remodeling to create longer and more branched arterial networks. This arterializing effect was not restricted to coronary vessels. *Dach1* overexpression also increased CX40-expressing artery branches in the retinal vasculature, and individual ECs overexpressing *Dach1* in the retina preferentially localized to arteries. ScRNAseq of *Dach1^OE^* coronary ECs revealed that it upregulated previously described arterial genes specifically in pre-artery and arterial populations. However, some transcriptional changes in all ECs were correlated with arterial differentiation, e.g. some genes that defined the artery clusters were up and lipid metabolism genes were down in all *Dach1^OE^* ECs. Finally, we found that overexpression of *Dach1* in the adult endothelium improved survival and heart function following MI. These data identify increased Dach1 as a new pathway by which to stimulate arteriogenesis and improve recovery after cardiac injury.

Significant work over the past two decades has provided information on how arterial ECs are specified during embryonic development (Corada et al., 2014). Using the developing dorsal aorta as a model, evidence in multiple species showed that VEGF-Notch signaling are critical for artery differentiation (Lawson et al., 2001; Lawson, Vogel, & Weinstein, 2002). Input from other signaling pathways (Wnt), downstream effectors (Erk, PI3K), and transcription factors (ETS, Fox) were subsequently shown to modulate artery differentiation in concert with VEGF and Notch (Corada et al., 2010; Hong, Peterson, Hong, & Peterson, 2006; Neal et al., 2019; Seo et al., 2006; Sissaoui et al., 2020; Wythe et al., 2013; You et al., 2005). Recent work has confirmed that several of these pathways also control artery specification in the coronary vasculature (Travisano et al., 2019; Su et al., 2018; Wang et al., 2017; Wu et al., 2012).

Here, we report a mechanism to robustly increase arterial EC specification and extend artery vessel coverage in hearts by overexpressing *Dach1*. An important feature of this transgenic model was that it displayed no gross defects in heart development or lethality. In contrast, genetic or pharmacological alterations in Notch or VEGF have strong effects on endothelial cell cycle and dramatically change the number of ECs in the heart (Fang et al., 2017; Pontes-Quero et al., 2019; Travisano et al., 2019; Wang et al., 2017; Wu et al., 2012). Thus, manipulation of Notch or VEGF make it difficult to conclude whether changes in artery growth are due to cell autonomous effects or secondary to dysregulated heart development. Furthermore, any therapeutic intervention aimed at boosting artery growth should not have other deleterious effects on the vasculature. Dach1’s specificity suggests it might make it a good candidate to explore for translational capabilities.

Our experiments in the retinal vasculature support previous data proposing that Dach1 supports artery development by potentiating EC migration against the direction of blood flow into growing arteries (Chang et al., 2017). In the retina, cells at the tip of the growing angiogenic front are selected as pre-artery cells and eventually change direction to migrate into developing arteries (Pitulescu et al., 2017; Red-Horse & Siekmann, 2019; Xu et al., 2014). By mosaically overexpressing *Dach1* in retinal ECs, we found that *Dach1^OE^* caused cells to occupy the tip cell position at early stages of vessel outgrowth. At later stages, *Dach1^OE^* cells moved from this tip position and were instead located in the mature artery. These data strongly support previous *in vitro* evidence that *Dach1* contributes to artery growth through driving flow directed EC migration (Chang et al., 2017). Future studies will utilize scRNAseq data to explore the mechanisms underlying this behavior.

Previously, we showed that increasing artery coverage and collateral artery development supports cardiac regeneration after MI (Das et al., 2019). Therefore, we hypothesized that overexpression of *Dach1* might produce the same effects during experimental cardiac injury. Animals with EC-specific *Dach1* overexpression experienced significantly increased ejection fraction and survival. Notable was significantly decreased fibrosis with a *Dach1^OE^*-specific pattern where damaged myocardium was restricted to an intramyocardial region and covered by surviving myocardium on both the epicardial and endocardial faces. This pattern was also evident in a recent study reporting enhanced cardiac regeneration following Meis1 and HoxB13 deletion (Uyen et al., 2020). It is tempting to speculate that vascular changes imparted by *Dach1^OE^* preserved these myocardial domains, and that this contributed to better function and survival.

In summary, we have discovered that overexpression of *Dach1* increases arterial EC specification in ECs during coronary and retinal development in mice. Dach1 did not act as a master regulator of artery cell fate but strongly enhanced artery cell fate commitment in specific sub-populations of ECs. In adults, recovery from MI was enhanced in *Dach1* overexpressing mice suggesting that promoting artery differentiation may be important therapeutically.

## Methods

### Animals

#### Mouse strains

Mouse husbandry and experimentation followed Stanford University Institution Animal Care and Use Committee (IACUC) guidelines. *ApjCreER* (H. I. Chen et al., 2014), *Cdh5CreER* (Sörensen et al., 2009), *CX40CreER* (Miquerol et al., 2015), *TdTomato* (The Jackson Laboratory, B6.Cg-Gt(ROSA)26Sortm9(CAG-tdTomato)Hze/J, Stock #007909), *Dach1^OE^* (see below), and CD1 (Charles River Laboratories, strain code: 022) mouse lines were used in this study.

The *Dach1^OE^* line was created by injecting a gRNA targeting intron 1 of the *ROSA26* locus, a repair template for the *Dach1^OE^* construct, and Cas9 mRNA into fertilized mouse eggs from the C57BL/6 mouse strain. The resulting F_0_ offspring were then genotyped by PCR for insertion of the transgene and propagated using C57BL/6 wild type mice. Predicted off target effects of the gRNA were made in silico, and those regions were sequenced to confirm that no erroneous cutting occurred at any of those sites. The *Dach1^OE^* repair template was generated by placing the CAG promoter upstream of a LoxP flanked 3x SV40 poly A region. This was followed by the Mouse CDS of *Dach1*, an IRES site, EGFP, and a final polyA sequence (Fig. 1A). The entire insert was flanked on both sides by homology arms to the *ROSA26* locus. For *Dach1^OE^* experiments, *Dach1^OE^* was crossed to either *ApjCreER, CX40CreER,* or *Cdh5CreER* which were maintained on mixed backgrounds. This breeding scheme generated Cre positive *Dach1^OE^*/ *Dach1^OE^* studs which were then crossed to wild type Cd1 (Jax) mice to generate either Cre^+^ *Dach1^OE^*/+ mice or Cre^−^ *Dach1^OE^*/+ littermates for controls.

#### Breeding and Tamoxifen Injection

Timed pregnancies were determined by defining the day on which a plug was found as E0.5. For Cre inductions, Tamoxifen (Sigma-Aldrich, T5648) was dissolved in corn oil at a concentration of 20 mg/ml and 4 mg was injected into the peritoneal cavities of pregnant dams (Fig. 1–2). For post-natal studies using the retinal vasculature P0 was defined as the day the mouse gave birth (Fig. 3). Tamoxifen was injected into the pregnant mother to allow the drug to transfer to the pups by lactation. For mosaic induction of *Dach1* in post-natal retinas Tamoxifen was diluted 1:100 in corn oil and .04 mg was given to the mother. For complete transgene recombination 4 mg was given to the mother. (Fig. 3) To induce *Dach1* overexpression in post-natal mouse hearts, 2 mg of Tamoxifen was injected at p0 to the mother (Fig. 2).

### Immunofluorescence and Imaging

#### Embryos

Embryos were dissected from the mother and fixed in 4% PFA for one hour at 4°C followed by three 15 minute washes in PBS. Hearts were then fully removed from the fixed embryo and placed in PBT (.5% Triton X-100 in PBS). Primary antibodies were then added to the embryos in PBT in 1.7 ml tubes and incubated overnight with rocking at 4°C. The following day, the hearts were washed every hour for 8 hours with rocking in PBT and then secondary antibodies were added in PBT and incubated overnight at 4°C with rocking followed by the same washing scheme. Finally, the tissue was transferred to vVectashield (Vector Laboratories, H-1000) and oriented with on the right, left, dorsal, or ventral side in a glass chamber slide. Images were then captured by confocal microscopy.

#### Retinas

Neonatal eyes were removed from the mouse and fixed in 4% PFA at 4°C followed by 3 × 15 minute PBS washes. Subsequently, the retina of each eye was dissected and four slits were cut down toward the center to make the retina flat for mounting later. Immunofluorescent staining was done using the same two day protocol as for the embryos with antibodies diluted in PBT in 1.7 ml tubes. To mount, retinas were placed on a glass slide and positioned with the four leaflets facing outward, followed by application of a thin layer of Vectashield, and finally the tissue was secured with a coverslip and nail polish.

#### Post-Natal Hearts

For post-natal hearts, samples were fixed in 4% PFA for 1 hour at 4 °C with rocking and washed twice for 15 minutes eash with PBS at 4 °C with rocking. The hearts were then incubated with primary antibodies diluted in PBT, and hearts were rocked at room temperature for 6 h and overnight at 4°C. To wash the primary antibodies, hearts were rocked in PBT at room temperature for 10 h and overnight at 4 °C. Hearts were washed in 50 ml PBT and the wash was changed every 2 h while rocking at room temperature. Hearts were then placed in secondary antibodies, diluted in PBT, at room temperature with rocking for 6 h and overnight with rocking at 4 °C. Hearts were then washed in 50 ml PBT for 8 h (wash changed every 2 h) and overnight at 4 °C. The washing was repeated for six more days. Prior to imaging, Vectashield was added to hearts in clean tubes, and hearts were equilibrated at room temperature for 40 min.

#### Immunohistochemistry

Tissue was fixed in 4% PFA for 1 hour at 4°C and washed 3 × 15 minutes in PBS. The samples were then dehydrated in 30% sucrose overnight at 4°C, transferred to OCT for a 2 hour incubation period, and frozen at −80°C. Cryosections through the tissue were made 20μm thick and captured on glass slides. Staining was performed by adding primary antibodies diluted in .5% PBT overnight and incubated with the sections at 4°C. The following day the slides were washed in PBS 3 times 10 minutes followed by 2 hour room temperature incubation with secondary antibodies, three more 10 minute washes, followed by mounting with Vectashield and a coverslip fastened using nail polish.

#### Microscopy and Image Processing

Images were captured on an Axioimager A2 epifluorescence microscope or a Zeiss LSM-700 confocal microscope. For each experiment, littermate Cre^+^ and Cre^−^ embryos or pups were stained and imaged together using the same laser settings. For each experiment, laser intensity was set to capture the dynamic range of the signal. For comparing immunostained embryos, z-stacks were created using the same number of steps for all samples. Images were captured using Zen (Carl Zeiss), Axio-Vision (Carl Zeiss), and processed using FIJI (NIH), Zen (Carl Zeiss), Photoshop (Adobe), and Illustrator (Adobe).

#### Antibodies

The following primary antibodies were used: Ve-Cadherin (1:125; BD Biosciences, 550548), anti-DACH1 (1:500; Proteintech 10914-1-AP), anti-ERG (1:500; Abcam, ab92513), anti-GFP (1:1000; abcam, ab13970), anti-CX40 (Alpha Diagnostic International, CX40A, 1:300), anti-Car4 (1:500; R&D, AF2414), Fabp4 (1:500; abcam, ab13979). Secondary antibodies were Alexa fluor-conjugated antibodies (488, 555, 633) from Life Technologies used at 1:250.

### Quantification

#### Artery Area

To calculate total area of CX40 staining on embryonic hearts, confocal z stacks of embryonic hearts were captured using the same heart orientation for all samples. Then, FIJI was used to make a z stack using the same number of slices for all images. Each stack was manually traced along all region of the image that had positive CX40 staining and the total area of the trace was then measured. While tracing, the researcher was blind to the genotype. The area of the heart used to tracing was also calculated by drawing a perimeter around the heart region to allow normalization of each measurement based on size. Finally, the percentage of that area occupied by the CX40 trace was calculated as a percent of the total heart area.

#### Vessel Width

Images were processed in the same way described above and the width of each vessel segment was measured by drawing a line perpendicular to the length of the primary coronary artery branch.

#### Vessel Length, Branchpoints, Endpoints

Confocal images of hearts at e17.5 and post-natal were traced along all CX40 artery segments using ImageJ. These traced artery segments were then imported to angiotool (Zudaire, Gambardella, Kurcz, & Vermeren, 2011) to measure total length of the network, number of branch points, and number of endopoints. All values were normalized to the total area of each heart.

#### Quantification of Artery Phenotype in Retinas

For each retina, one randomly chosen quarter leaflet was used for quantification. Overlapping images were taken of the retina and then stitched using FIJI. The total length of all CX40 positive segments were then manually traced for each quarter retina for both control and *Dach1^OE^* retinas with the genotype blinded to the investigator. The cumulative distance of each CX40 trace was then compared between the two genotypes. To determine artery and vein crossings, arteries were defined by CX40 positive vessels, and veins were CX40 negative. Each crossing per quarter retina was counted and compared between control and *Dach1^OE^* retinas.

#### Mosaic Analysis of Retinas

To analyze the distribution of all labeled cells, each cell was counted and given a label based on whether it was located in the capillary, tip, vein, or artery. Tip cells were classified as any cell on the outermost edge of the vascular front. Arteries and veins were identified based on morphology with arteries as thinner vessels and aligned cells, and veins with thicker vessels and more rounded cells. Capillaries were all other cells not at the tip and with diameter of one cell width. The total number of cells in each compartment was then summed and the percent of cells in each compartment was graphed.

#### Statistical Analysis

Unpaired t tests were used to determine the two tailed P-value for each comparison of two groups (i.e. *Dach1^OE^* v Control). Two-way ANOVA was used when making comparisons across multiple samples. A Logrank test was used to determine the statistical significance of the survival curve. Prism 8 was used to generate graphs and perform statistical analysis.

### Single Cell RNA seq

*ApjCreER;Dach1^OE^/TdTomato* male mice were crossed to female Cd1 wild type mice to generate *ApjCreER;Dach1^OE^/+* or *ApjCreER;TdTomato*/+ sibling matched embryos. To generate enough samples for the experiment, approximately 8 litters were used totaling 10-15 embryos per group. The hearts from all embryos were then sorted into two tubes based on fluoresce signal and digested with 300μl 500 U/ml collagenase IV (Worthington #LS004186), 1.2 U/ml dispase (Worthington #LS02100), 32 U/ml DNase I (Worthington #LS002007), and sterile DPBS with Mg2+ and Ca2+ at 37°C for 45 minutes with pipet mixing every 7 minutes. Once digestion was complete, 60μl FBS and 1.2 ml PBS were added to the suspension and filtered with a 40 μm strainer. After washing once in 3% FBS in PBS, FACS antibodies were added at a concentration of 1:20 and incubated on ice for 40 minutes. Cells were then washed once more in 3% FBS in PBS and resuspended in 1 ml of 3% FBS for FACS. The following FACS antibodies were used: APC-CD31 (eBiosciences, 12.5ul/ml), FITC-CD31 (eBiosciences, 12.5ul/ml), APC-Cy7 CD45 (Biolegend 12.5ul/ml), APC Cy7-Ter119 (Biolegend 12.5ul/ml) and Dapi. Once stained, the cells were sorted on a BD FACS Aria II FACS machine into 1.7 ml tubes. The gates were set up to sort cells with low DAPI, high CD31 (endothelial marker), high EGFP (*Dach^OE^*) or high Td (control), low CD45 (hematopoietic cells), and low Ter119 (erythroid cells). Compensation controls were set up for each single channel (EGFP, TdTomato, Dapi, CD31, and combined CD45 and Ter119) before sorting the final cells.

*Dach1^OE^* and control coronary ECs were then submitted to the Stanford Genome Sequencing Service Center for 10X single cell V3 library preparation. Sequencing was done using Illumina HiSeq 4000.

Initial processing of raw Illumina reads was performed using Cell Ranger (10X Genomics). Raw BCL files were demultiplexed and converted to FASTQ files using the mkfastq function. Subsequently, reads were aligned to the mouse genome (mm10) as well as *EGFP* and *tdTomato* sequences using the count function. Sequencing and alignment statistics are shown in Supplemental Figure 2.

The majority of scRNA-Seq data analysis was performed using R and Seurat. Cells were deemed low-quality and excluded from downstream analysis if they 1) expressed more than 6500 genes (doublets) 2) expressed less than 2000 genes (dead cells) or 3) if more than 10% of counts aligned to mitochondrial genes (dead cells). These cutoffs were determined from the distribution of genes expressed and mitochondrial transcripts (Supplemental Figure 1). We also removed contaminating tdTomato+ cells from the *Dach1^OE^* group, EGFP+ cells from the control group, and a cluster of Gpr126+ endocardial cells from both groups (Supplemental Figure 2). In total, 1975 of the initial 3294 control cells and 2149 of the initial 3368 *Dach1^OE^* cells were deemed high-quality ECs suitable for analysis.

We first generated individual Seurat objects for control and *Dach1^OE^* ECs. Following data normalization and variable feature selection, control and *Dach1^OE^* datasets were integrated (Stuart et al., 2019). To reduce the dimensionality of the dataset, we performed principal component analysis on the shared variable features determined during integration. The top 50 principal components were then used for visualization and clustering. Of note, we did not observe any differences in clustering or visualization when using anywhere from 10 to 100 principal components for downstream analysis. To visualize scRNA-Seq data in two-dimensional space, we used the Uniform Manifold Approximation and Projection algorithm (McInnes *arXiv* 2018). We then identified subpopulations of ECs (i.e. clusters) by constructing a shared nearest neighbor graph (k=20) and detecting communities with the Louvain algorithm (Stuart et al., 2019). Transcripts enriched in each cluster compared to all other clusters were determined using the Wilcoxon Rank Sum test. These lists of enriched transcripts were then used to annotate EC subtypes. We performed this process of clustering, marker determination, and annotation with a range of inputs to the “resolution” parameter of the Louvain community detection algorithm. We found that a resolution of 0.7 gave clusters which could be most readily identified as EC subtypes. Gene expression differences between control and *Dach1^OE^* in various EC subpopulations were determined using the Wilcoxon Rank Sum Test. Artery, Vein, Lipid oxidation, Lipid transport, Lipid synthesis, and Eicosanoid metabolism scores were assigned to each cell by calculating the enrichment (Tirosh et al., 2016) of transcripts for each process or cellular state. We used markers of the “Artery 2” and “Vein” clusters as artery and vein-specific genes, respectively. Gene lists used to score enrichment of lipid-related processes were gathered from the Gene Ontology database (http://geneontology.org). To infer potential developmental relationships between EC subtypes, we performed trajectory analysis using Monocle 3 (Cao *Nature* 2019). Cellular UMAP embeddings were first imported from Seurat objects into Cell Data Set objects. Subsequently, we performed Louvain clustering with k=100 followed by reverse graph embedding for trajectory construction. CytoTRACE analysis (Gulati *Science 2020*) was performed as described at https://cytotrace.stanford.edu/.

ScRNA-Seq datasets from coronary ECs at e12.5 and e14.5 (Su et al., 2018) were analyzed to determine the relative distribution of EC subpopulations as a function of development. Datasets were integrated using the batch correction technique described by Stuart et al. To specifically isolate coronary ECs from the e12.5 dataset, sinus venosus and valvular cells were excluded.

We found genes that were up or downregulated by Dach1 in each cluster using a logFC > 0.25 and an adj. p value <0.05. DEGs in the four cycling capillary clusters were combined for subsequent analysis due to the similarity in these clusters. We then compared the DEG lists to the genes which were positively or negatively enriched in the control mature arterial EC cluster (Art 2). A smaller list of genes that were up or downregulated by Dach1 in all clusters was also used to find the overlap to control artery cluster genes. In addition, we generated a longer list of DEGs that were changed by Dach1 in any cluster. This list was used to compare to a list of genes regulated by Dach1 in vitro (Chang et al., 2017) or a list of genes correlated with *Dach1* expression in control hearts. To generate this list of *Dach1* correlated genes, we calculated the Pearson’s correlation coefficient between *Dach1* and all other transcripts expressed by control cells. The top 500 positively or negatively correlated genes were used for analysis, excluding ribosomal transcripts.

### Coronary Artery Ligation Experiments

#### Surgery

All MIs were performed by the same surgeon, who was blinded to genotype. *ApjCreER* mice were bred to *Dach1^OE^/Dach1^OE^* mice to generate either *ApjCreER;Dach1^OE^*/+ (*Dach1^OE^*) or *Dach1^OE^*/+ (control) mice. Once the mice reached 6 weeks of age, a single dose of 4 mg Tamoxifen was given. 6 weeks later, mice were subjected to permanent coronary artery ligation, under anesthesia using isofluorane. The chest cavity was opened and a 7-0 silk suture was placed around the left anterior descending artery (LAD), with occlusion verified by blanching of the underlying myocardium. The chest was then sutured closed. Following surgery, Buprenorphine (0.1 mg/kg) was used as an analgesic.

#### Assessments of cardiac function

Mice were monitored for survival for 4 weeks after the surgery. Transthoracic echocardiograms were performed before and after surgery under anesthesia with 1.5% isoflurane. At the completion of the study at 4 weeks, mice were sacrificed and hearts were collected in 4% PFA. Following fixation, ligated hearts were embedded in paraffin and sectioned at 10μM. Slides were stained with Masson’s Trichrome (Sigma, HT15-1KT) and imaged using a slide scanner. The experiment was performed twice, and from these a total of 30 control and 32 *Dach1^OE^* animals were included. A small number of animals were excluded from both the control and experimental groups if histological analysis showed no evidence of myocardial scarring using Masson’s Trichrome staining, which indicated an unsuccessful ligation.

#### Histology quantification

Scar area was measured in comparable sections by selecting the entire region covered by Masson’s Trichrome staining (i.e. blue staining) and expressing that number as a percentage of the total heart area calculated for the same section. To classify fibrosis patterns as either full thickness or mid-myocardial, any section where the fibrosis was bordered on the interior and/or exterior side by Masson’s Trichrome-negative heart muscle was considered mid-myocardial. Sections where the fibrosis was not bordered on the interior and exterior side by any tissue were counted as full thickness. To measure the width of the myocardium hearts were converted to binary images. Then, a point in the center of the left ventricle was positioned, and from this point, 360 radii were sequentially drawn. Along each of these radii, the length of black pixels along the line was measured and transformed into a graph where the thickness is visualized at all points around the heart. The zero angle corresponds to the center of the scarred area.

## Supporting information

Supplemental table 1 control art2 markers down

Supplemental table 1 control art2 markers up

## Acknowledgements

K.R. is a New York Stem Cell Foundation – Robertson Investigator. K.R. is also supported by the NIH (R01-HL128503). B.R. is supported by NIH (5 F31 HL147410-02), NSF GRFP. I.W. is supported by T32HL098049. P.R. is supported by NIH (T32GM007276), NSF GRFP. R.P. is supported by an AHA graduate fellowship. A Stanford CVI seed grant funded MI experiments. D.B is supported by the Department of Defense CMDRP in Congenital Heart Disease W81XWH-16-1-0727. We thank all members of the Red-Horse lab for technical and intellectual support. We thank member of the Stanford Genome Sequencing Services Center which is supported by NIH Grant # 1S10OD020141-01.

## Contributions

B.R., I.W., D.B., and K.R conceptualized the study and wrote the manuscript. B.R., I.W., P.R., and H.Y.Z. performed experiments and computational analysis. B.R., M.Z., and R.R. performed cardiac injury studies. A.C. designed the *Dach1^OE^* mouse. R.P and G.D. assisted with experimental procedures. K.R. and D.B. provided recourses.

---

**Supplementary Figure 1.**
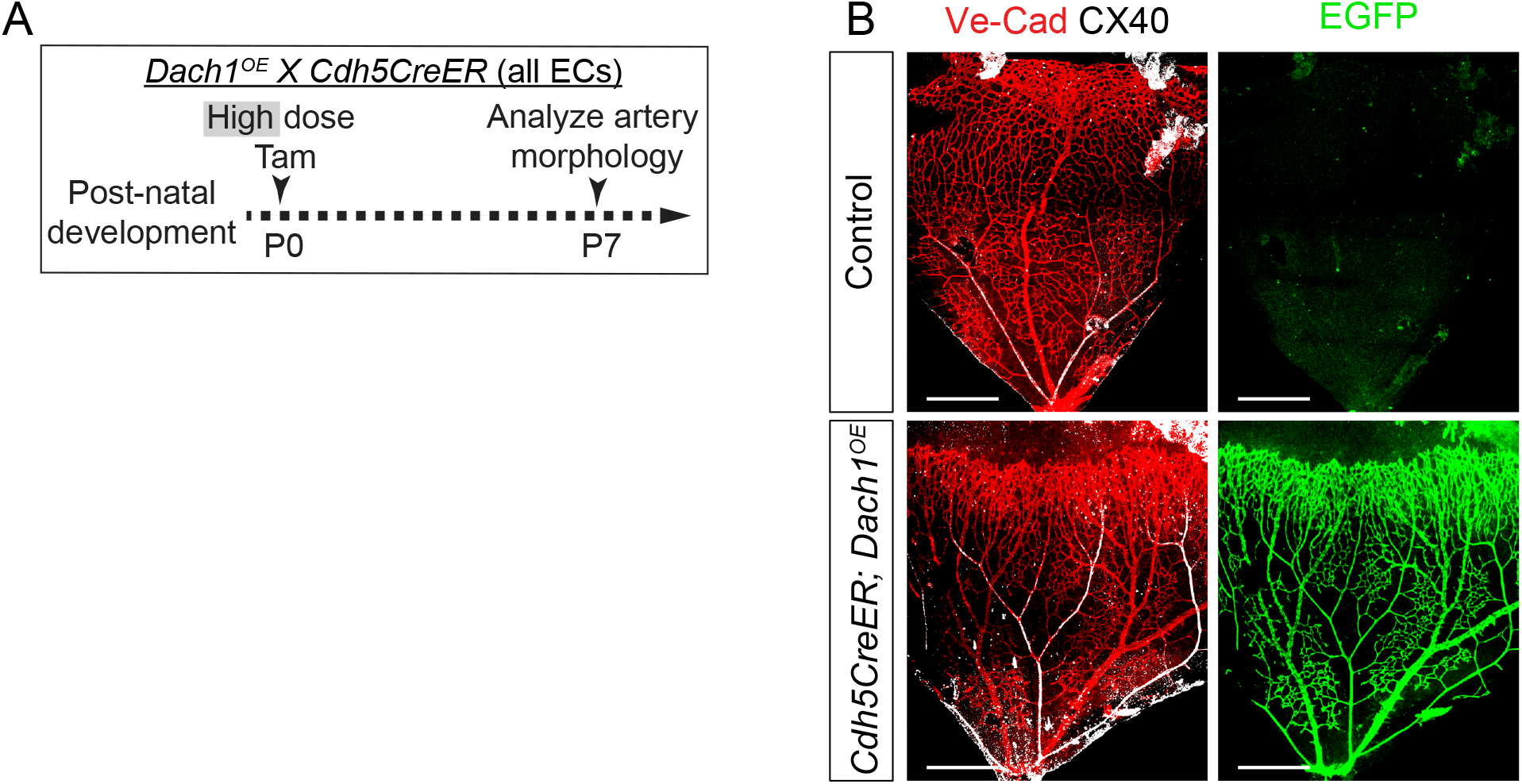
Recombination of *Dach1^OE^* in the retina. **A)** *Dach1^OE^* was crossed to *Cdh5CreER*, and pups were given Tamoxifen at P0 through lactation from the mother. **B)** At P7, retinas were stained for VE-cadherin, CX40, and EGFP to detect recombination of the *Dach1^OE^* allele. Scale bar= 400μM.

**Supplementary Figure 2.**
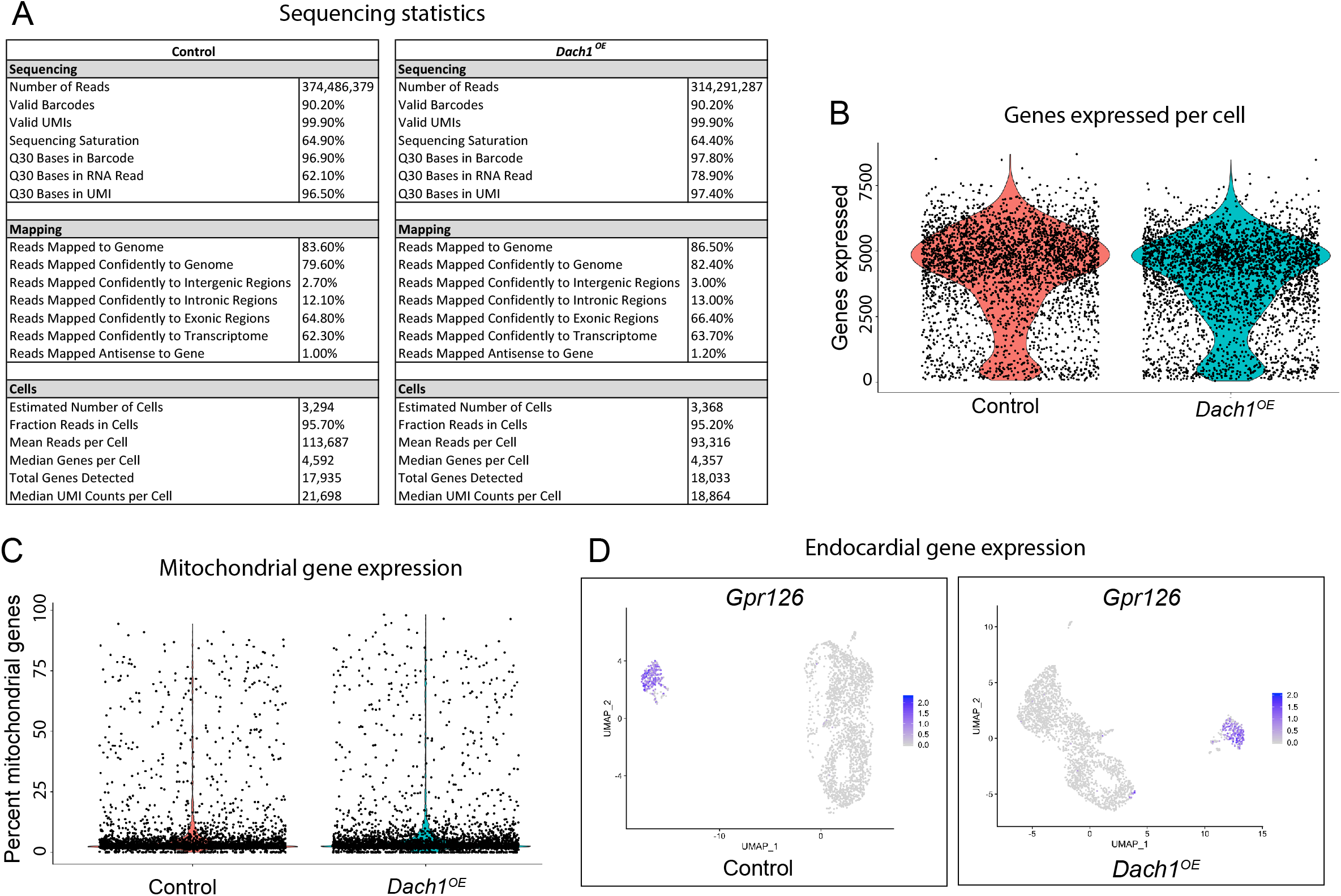
Quality control for single cell sequencing. **A)** Sequencing statistics from each group. **B)** Number of genes expressed per cell in both control (*TdTomato*) and *Dach1^OE^* hearts. Cells expressing more than 6500 and fewer than 200 were excluded for analysis. **C)** The percent of expressed genes mapping to the mitochondrial genome, cells with greater than 10% were excluded. **D)** *Gpr126* expression in control and *Dach1^OE^* samples. Cells expressing this endocardial marker were excluded.

**Supplementary Figure 3.**
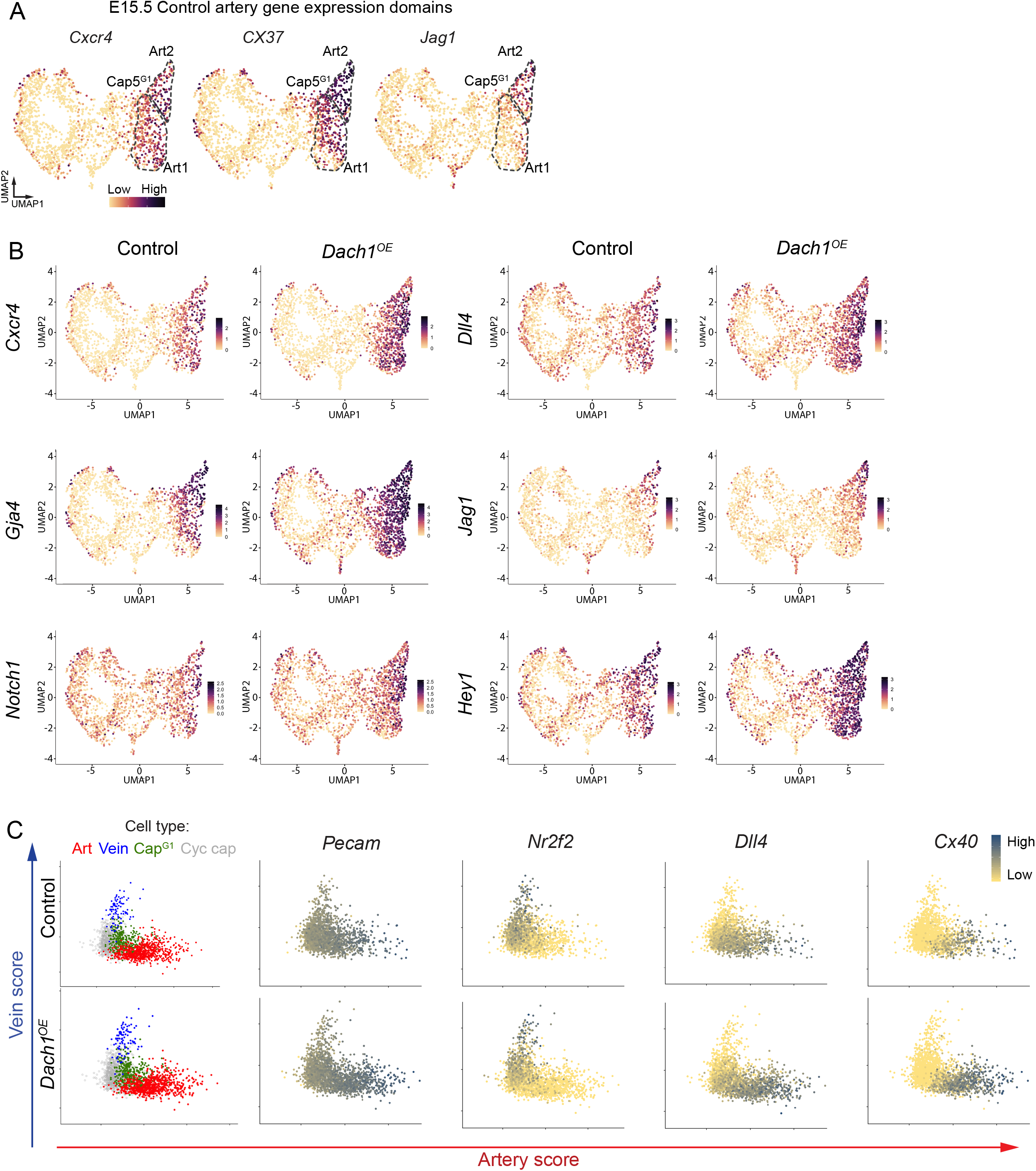
Artery expression continuum. **A)** Expression of three artery genes plotted on UMAP plots, occupying different domains between Art1 and Art2. **B)** The expression pattern of a set of canonical artery markers for control and *Dach1^OE^* cardiac endothelial cells. *Dach1* overexpression expands the expression of some but not all of the markers. **C)** Each cell in the scRNA seq data set was scored based on its cumulative expression of all artery 2 cluster marker genes and its expression of all venous cluster genes. Each cell was then plotted with the x axis as the artery score and the y axis as the vein score. Cells from *Dach1^OE^* hearts are shifted along this continuum to have greater artery gene expression.

**Supplementary Figure 4.**
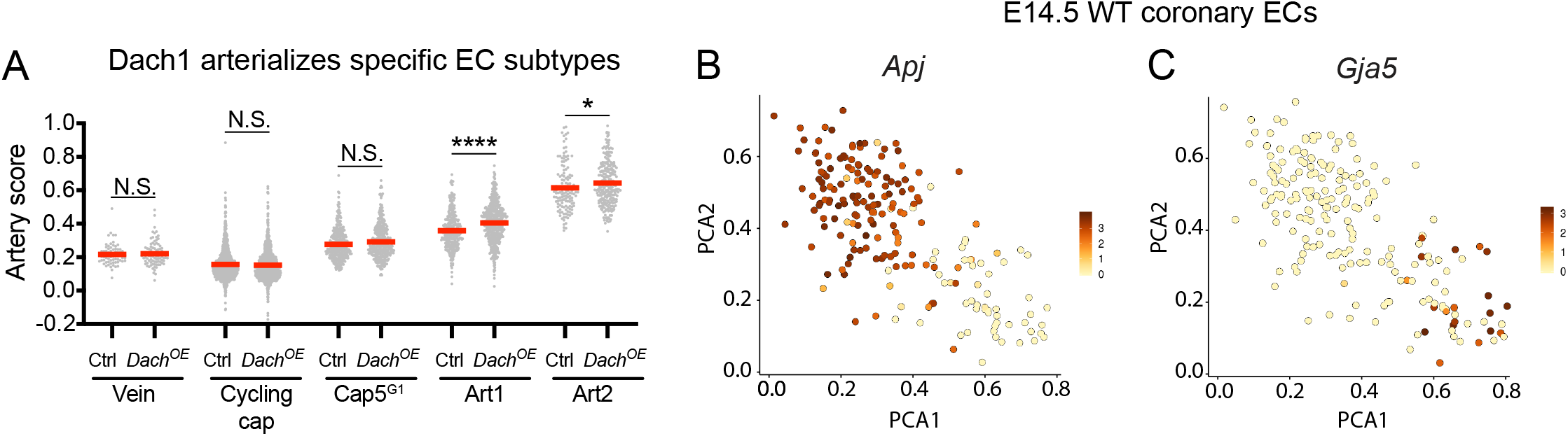
Artery lineage pathway. **A)** Artery score of each sample for every cluster. **B,C)** PCA plot of wild type cardiac endothelial cells at e14.5 from Su et al. **B)** *Apj* is expressed in all endothelial cell types except the artery at this stage. **C)** Location of artery cells (*Gja5*^+^) on the PCA plot.

**Supplemental Figure 5.**
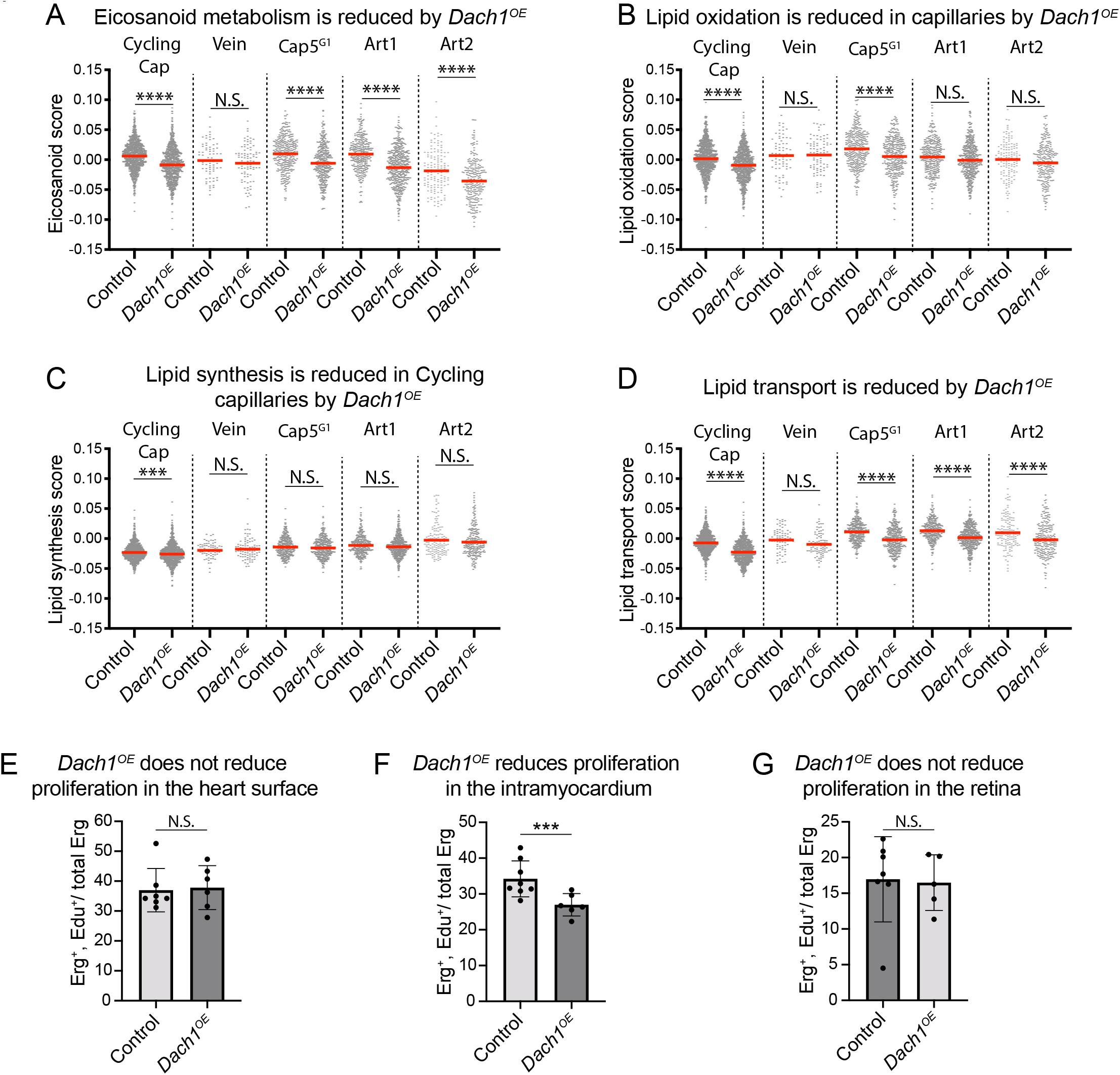
Lipid and proliferation pathways downstream of Dach1. **A-D)** Each cell was scored based on its expression of genes in different lipid pathways. Across pathways and clusters, *Dach1^OE^* tended to decrease scores. **E, F)** E15.5 embryos from Cre^−^ and *Cdh5CreER, Dach1^OE^* littermates were dosed with Edu by IP injection to the mother to assess the proliferation rate of endothelial cells. Regions of interest in either the surface or intramyocardial region of the capillary plexus were selected for measurement. The Edu/Erg ratio was reduced by *Dach1^OE^* only in the intramyocardial region (n=7 Cre^−^, n=6 *Cdh5CreER, Dach1^OE^*). **G)** P6 mice from control and *Cdh5CreER; Dach1^OE^* were dosed at P0 with Tamoxifen nd then at P6 with Edu. Retinas were then dissected and stained for Erg and Edu. There was no significant difference in proliferation between control and *Cdh5CreER; Dach1^OE^* retinas (N=7 control, n=5 *Dach1^OE^*). N.S.= not significant, ***=p<.001, ****=p<.0001, red bar= mean, all data are mean+-SD.

**Supplemental Table 1.** Lists of all genes that define the artery 2 cluster in control hearts. “Artery genes” are upregulated cluster markers, while “Non-artery genes” are downregulated cluster markers.

## Notes

### Competing Interest Statement

The authors have declared no competing interest.

